# Hypothermic and Normothermic Machine Perfusion Differentially Shape the Renal Cellular Landscape

**DOI:** 10.64898/2026.10.07.757234

**Authors:** Janine Siegle, Dörte-Maria Kemper, Runze Zhu, Yiren Yang, Jay Wehenkel, Gishnu Harikumar Parvathy, Riccardo Camiletti, Karolin Hublitz, Efstathios G. Stamatiades, Vanessa Pross, Robert Labes, Michael Fähling, Roman Bülow, Peter Boor, Katja Kotsch

**Affiliations:** Department of General- and Visceral Surgery, Charité – Universitätsmedizin Berlin, Campus Benjamin Franklin, corporate member of Freie Universität Berlin and Humboldt-Universität zu Berlin, Germany; Department of Microbiology, Infectious Diseases and Immunology, Charité – Universitätsmedizin Berlin, Campus Benjamin Franklin, corporate member of Freie Universität Berlin and Humboldt-Universität zu Berlin, Germany; Institute of Translational Physiology (CCM), Charité – Universitätsmedizin Berlin, corporate member of Freie Universität Berlin and Humboldt-Universität zu Berlin, Germany; Institute of Pathology, University Clinic Aachen, RWTH Aachen University, Aachen, Germany

## Abstract

The shortage of donor organs has led to an increased use of extended-criteria donors and donation after circulatory death for kidney transplantation, both of which are associated with graft injury. Although machine perfusion (MP) offers a preservation strategy, the most beneficial approach for kidneys with respect to effective graft preservation remains unclear. We compared oxygenated hypothermic (HMP) and normothermic machine perfusion (NMP) using a murine kidney perfusion model regarding intra-graft immune cell composition, viability, and inflammatory responses. NMP resulted in more severe tubular injury, accompanied by increased immune cell mobilization out of the graft. NMP increased early apoptosis among kidney-resident immune-cell populations, whereas endothelial cells showed increased late apoptosis after both strategies. Proximal tubular-cell apoptosis remained comparable to naïve tissue, and neither modality reduced reactive oxygen species levels. Both protocols induced antioxidant and hypoxia-responsive genes compared with static cold storage, while NMP elicited a stronger inflammatory response. Treatment with the senolytic ABT-263 demonstrated the model’s feasibility for investigating renal injury, cellular viability, and immune activation during MP.

## Introduction

The implementation of machine perfusion (MP) is transforming the field of solid organ transplantation, particularly by expanding the utilization of marginal donor organs. While normothermic machine perfusion (NMP) has successfully increased the use of transplantable livers, comparable advances in kidney transplantation remain limited^1–3^. Hypothermic machine perfusion (HMP), which preserves kidneys in a metabolically quiescent state, has become the standard preservation strategy because of its superior outcomes compared with static cold storage (SCS) in improving both early and long-term graft outcomes^4,5^. In contrast, NMP maintains perfused organs under near-physiological conditions, enabling functional viability assessment before transplantation but also serving as a platform for targeted therapeutic interventions. By restoring oxidative metabolism, NMP reduces susceptibility to mitochondrial dysfunction^6,7^. Consistent with this, porcine kidneys preserved by NMP exhibit improved mitochondrial integrity and higher adenosine triphosphate (ATP) levels than kidneys subjected to SCS. However, restoring cellular metabolism during NMP may also increase metabolic injury. Further investigation is therefore needed to clarify the mechanisms underlying these effects^8,9^.

Despite considerable advances in our understanding of renal metabolism during MP, the effects of different perfusion strategies on renal tissue integrity and the graft-resident immune compartment remain unclear. Kidney-resident leukocytes, including tissue-resident macrophages, contribute to immune surveillance as well as tissue homeostasis, metabolic adaptation, and repair following transplantation^10^. However, whether hypothermic and normothermic perfusion preserve these populations to a similar extent is unknown. The same applies to structural cell populations, including endothelial cells (ECs) and proximal tubular epithelial cells (PTECs). This question is particularly relevant for graft-directed therapies administered during MP, including siRNA^11^, mesenchymal stromal cells^12^, and receptor antagonists^13^. To date, the efficacy of these approaches has been assessed mainly on the basis of functional outcomes, with less attention to their effects on renal cellular composition and tissue integrity. Building on our previous observation that donor kidneys retain distinct donor-specific cellular signatures^14^, we investigated whether normothermic and hypothermic MP differentially affect the intrarenal cellular landscape. We focused on graft-resident leukocyte composition, apoptosis and necrosis, and the preservation of structural cell populations, including ECs and PTECs. Together, these analyses provide a cellular perspective on how MP strategies affect the intrarenal microenvironment.

## Results

### NMP increases vascular resistance and accelerates tubular injury

To assess renal integrity following HMP and NMP, we used a murine kidney perfusion model that enables oxygenated perfusion at either 4°C (HMP) or 37°C (NMP)^15^. The experimental design is shown in Figures 1A and 1B. Both perfusion modalities supported stable *ex vivo* kidney perfusion for 60 minutes, with vascular resistance remaining below 50 mmHg/mL/min throughout the experiments. Vascular resistance differed between the perfusion conditions, with the highest observed during NMP supplemented with 30% red blood cells (RBCs) compared with NMP supplemented with 15% RBCs and of HMP (Figure 1C). Flow rates were also significantly higher during HMP than during NMP with 30% RBC supplementation (Figure 1D). After 60 minutes of perfusion, free hemoglobin levels were significantly lower with 15% than 30% RBCs during NMP (Figure 1E), and creatinine clearance was significantly higher, indicating better glomerular function (Figure 1F). In contrast, fractional sodium excretion did not differ significantly between groups, indicating comparable tubular function (Figure 1G). However, histological analysis revealed more acute tubular injury after NMP than after HMP, even with 15% RBCs. Therefore, subsequent analyses throughout the study were performed using 15% RBCs (Figures 1H and 1I). Gene expression analysis showed significantly lower kidney injury molecule-1 (KIM-1; *Havcr1*) mRNA levels in SCS kidneys than in naïve controls, with no differences among SCS, HMP, and NMP. In contrast, lipocalin-2 (NGAL; *Lcn2*) expression remained comparable across all groups (Figure 1J).

**Figure 1.**
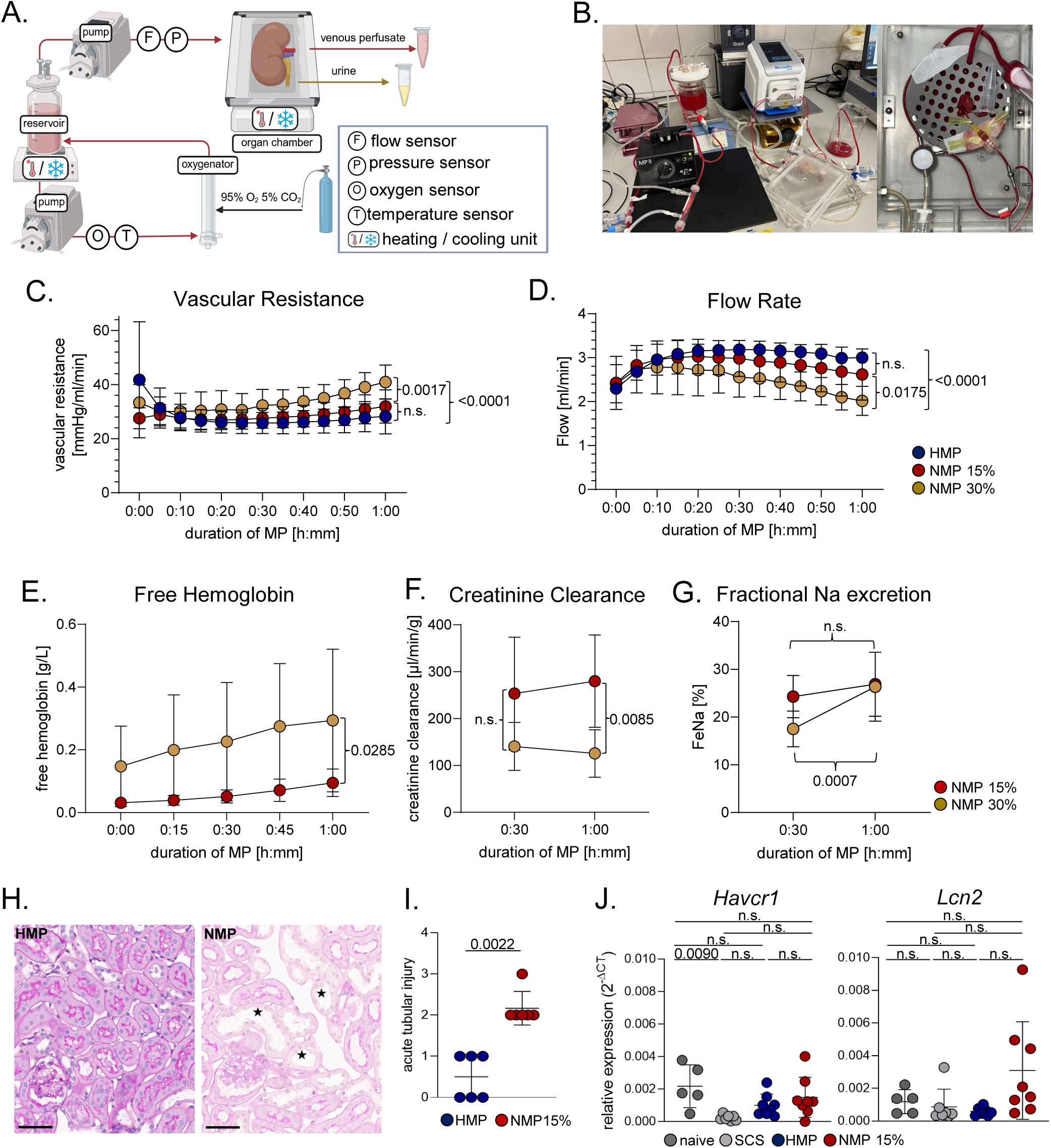
Hypothermic (HMP) and normothermic machine perfusion (NMP) of murine kidneys. (A) Schematic overview of the machine perfusion (MP) setup used for oxygenated HMP and NMP. (B) Experimental setup used for oxygenated NMP, including the perfusion circuit and kidney chamber. (C) Renal vascular resistance (VasRes) and (D) renal blood flow during 1 h of HMP and NMP, with NMP performed using perfusate supplemented with either 15% or 30% erythrocytes. Data points are mean ± standard deviation (SD), n=13/experimental group. Statistical significance was determined using a two-way ANOVA with Tukey’s post-hoc test. (E) free hemoglobin, (F) Creatinine clearance and (G) Fractional Na^+^ excretion during NMP using perfusate supplemented with either 15% or 30% erythrocytes. Data points are mean ± SD, n=5-8/experimental group. Statistical significance was determined using a mixed-effects analysis with Sidak’s multiple comparisons test. (H) Representative images and (I) quantitative analysis of the histopathological evaluation of HMP- and NMP-perfused kidneys. Asterisks indicate tubular dilatation as a form of acute tubular injury, scale bar: 60 µm. Individual data points are shown with mean ± SD, n=6/experimental group. Statistical significance was determined using Mann–Whitney *U* test. (J) Gene expression analysis of *Havcr1* (kidney injury molecule-1, KIM-1) and *Lcn2* (lipocalin-2, NGAL) in naïve kidneys, kidneys subjected to 1 h of static cold storage (SCS), and kidneys undergoing HMP or NMP. Individual data points are shown with mean ± SD, n=5-8/experimental group. Statistical significance was determined using two-way ANOVA with Tukey’s multiple comparisons test.

### Immune cell composition of the perfusate is shaped by the perfusion protocol

During MP, leukocytes exit the organ via diapedesis^16,17^. To assess whether the perfusion protocol has an impact on immune cell populations released from the graft, we analyzed the cellular composition of perfusates following 1 hour of HMP or NMP (Figure 2A). Optimized t-Distributed Stochastic Neighbor Embedding (Opt-SNE) analysis revealed distinct immune cell clusters associated with each perfusion modality (Figure 2B). Compared with HMP, NMP was characterized by significantly lower frequencies of CD4⁺ T cells, CD4⁺CD8⁺ double-positive (DP) T cells, B220^+^ B cells, F4/80^+^CD11b^+^ macrophages, and MHCII⁺CD11c⁺ dendritic cells in the perfusate, whereas Ly6G⁺CD11b⁺ neutrophils were preferentially released during NMP. No differences were detected for CD8^+^ T cells, CD25^+^FoxP3^+^ regulatory T (Treg) cells or SigF^+^CD11b^+^ eosinophils Figure (2C, D). We further examined various T cell subsets and found a preferential release of CD4⁺CD44⁺CD62L⁻ memory T cells during NMP into the perfusate. Interestingly, this difference was not evident within the CD8⁺ T cell compartment (Figure 2E). Conversely, HMP perfusates contained higher frequencies of naïve CD8^+^ T cells (Figure 2E). No differences were observed between HMP and NMP for CD4^+^ and CD8^+^ tissue-resident cells identified by their CD69 and CD103 expression (Figure 2E).

**Figure 2.**
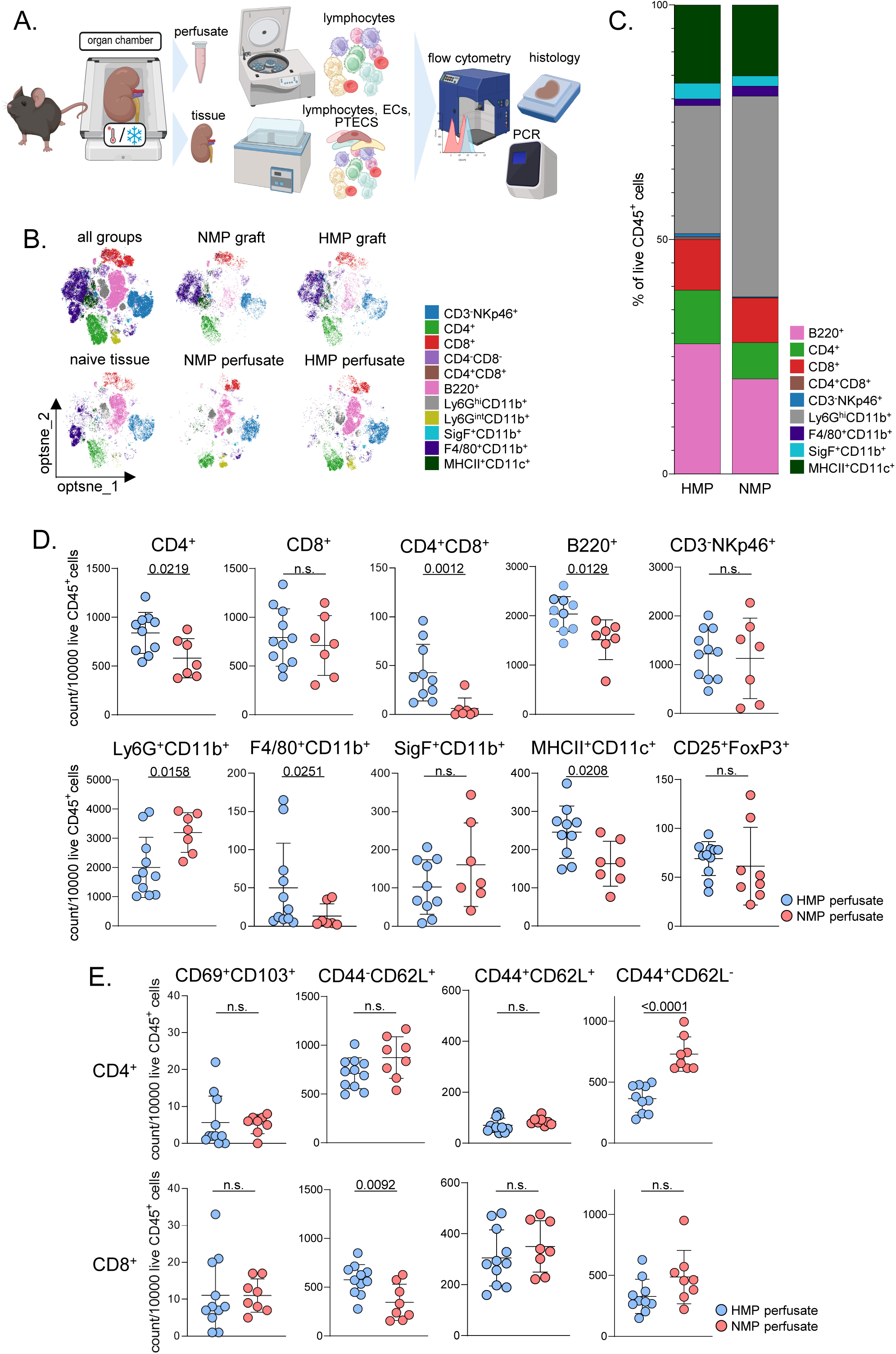
Leukocytes released into the perfusate during HMP and NMP. (A) Schematic overview of the experimental workflow. After HMP and NMP, lymphocytes were enriched from the perfusate. In parallel, leukocytes, glomerular endothelial cells (ECs), and proximal tubular epithelial cells (PTECs) were isolated from the corresponding perfused kidneys for analysis by flow cytometry, histology, and real-time RT-PCR. (B) Opt-SNE plots illustrating the distribution of leukocyte populations in naïve grafts and grafts following HMP or NMP perfusion. Cells were gated on live CD45⁺ cells (n=11/experimental group). (C) Proportion of live CD45⁺ leukocytes in perfusates collected from HMP- and NMP-perfused grafts (n=11/experimental group). Data shown are means scaled up to 100%. (D) Cell counts of the major leukocyte populations identified in HMP and NMP perfusates. (E) Cell counts of CD4^+^ and CD8^+^ tissue-resident T cells (CD69⁺CD103⁺), as well as naïve (CD44⁻CD62L⁺), central memory (CD44⁺CD62L⁺), and effector memory (CD44⁺CD62L⁻) T-cell subsets (n=7-10). Individual data points are shown with mean ± SD; n=7-11/experimental group, some samples were excluded due to live CD45^+^ cell numbers < 10,000 cells. Data distribution was assessed using the Kolmogorov–Smirnov test. Statistical significance was determined using Welch’s *t*-test or Mann–Whitney *U* test, as appropriate.

### Machine perfusion alters the composition of graft-resident immune cells

Next, we compared immune cell subsets within HMP-and NMP-perfused grafts with those detected in the corresponding perfusates. As shown in Figure 3A, naïve kidneys contained significantly higher frequencies of B cells than perfused grafts, consistent with their mobilization into the perfusate during MP. CD4⁺ T cells were preferentially detected within the kidney during NMP, whereas CD8⁺ T cell frequencies did not differ between grafts and perfusates under either condition (Fig. 3B, C). Similarly, DP T cells, antigen-presenting macrophages, and dendritic cells were preferentially retained within the graft. In contrast, natural killer (NK) cells were significantly released into the perfusate under HMP, while neutrophils were efficiently mobilized under both conditions. Eosinophils were detected at higher frequencies in NMP perfusates, and Tregs showed a trend toward release under HMP (Figure 3D). Further characterization of T cell subsets showed mobilization of naïve CD4⁺ and CD8⁺ T cells into the perfusate under both conditions. Similarly, CD44⁺CD62L⁺ central memory T cells were mobilized, whereas CD44⁺CD62L⁻ effector memory T cells were preferentially retained within the graft. Tissue-resident CD103⁺CD69⁺ CD4⁺ and CD8⁺ T cells were likewise retained, suggesting preferential preservation of tissue-resident immune populations during NMP (Figure 3E). Overall, NMP and HMP differentially affected immune-cell trafficking, highlighting perfusion modality as a determinant of intra-graft immune dynamics during *ex vivo* preservation.

**Figure 3.**
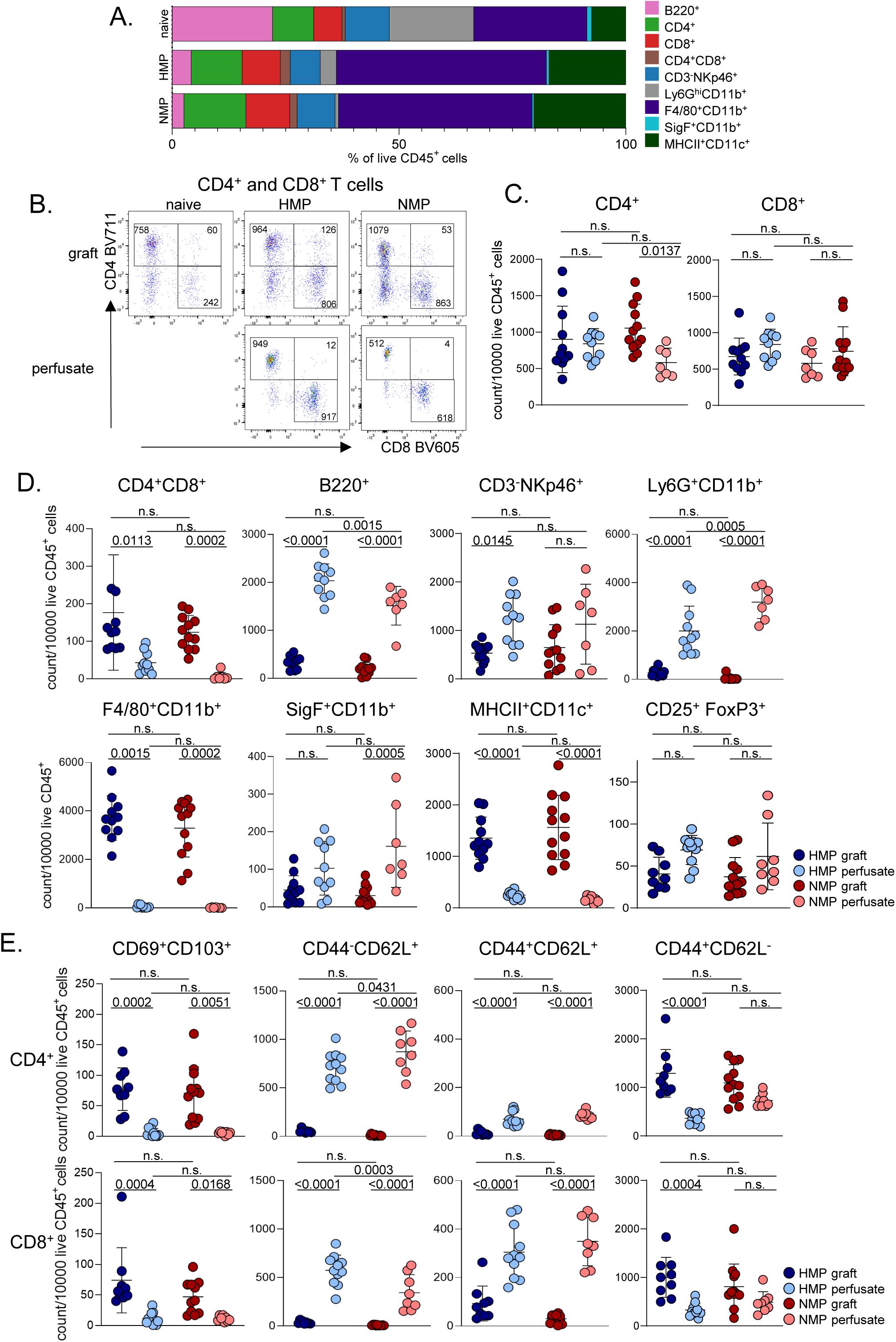
HMP and NMP differentially shape the intra-graft leukocyte composition. (A) Bar graphs show the relative abundance of the major renal leukocyte populations in HMP- and NMP-perfused kidneys compared with naïve control kidneys. (n=11/experimental group). Data shown are means scaled up to 100%. (B) Representative flow cytometry dot plots showing the cell numbers/10,000 live CD45^+^ cells of CD4^+^, CD8^+^ and CD4^+^ CD8^+^ T cells isolated from naïve tissue, perfused grafts and perfusates. (C) Comparison of the cell counts of CD4^+^ and CD8^+^ T cells and (D) comparison of the cell counts of the major leukocyte populations identified in HMP and NMP perfusates and grafts. (E) Comparison of cell counts of CD4^+^ and CD8^+^ tissue-resident T cells (CD69⁺CD103⁺), as well as naïve (CD44⁻CD62L⁺), central memory (CD44⁺CD62L⁺), and effector memory (CD44⁺CD62L⁻) T-cell subsets identified in HMP and NMP perfusates and grafts. Individual data points are shown with mean ± SD; n=7-11/experimental group. Data distribution was assessed using the Kolmogorov–Smirnov test. Statistical significance was determined using ordinary one-way ANOVA with Tukey’s multiple comparisons test or Kruskal-Wallis test, as appropriate.

### NMP enhances early apoptosis cell death

We further investigated the effects of HMP and NMP on immune cell viability in the graft and in the perfusate by assessing apoptosis and necrosis. Using Annexin V and L/D Zombie aqua staining, immune cells were classified as viable (LIVE, Zombie^-^Annexin V^-^), early apoptotic (EA, Zombie^-^Annexin V^+^), late apoptotic (LA, Zombie^+^Annexin V^+^), or necrotic (N, Zombie^+^Annexin V^-^), as exemplified for CD4⁺ T cells in Figure 4A. Among the non-viable cells derived from the graft, the most abundant population was early apoptotic F4/80⁺ macrophages, whereas neutrophils and eosinophils were detected at the lowest frequencies. By contrast, the most prominent non-viable populations in the perfusate were B cells, NK cells and neutrophils (Figure 4B). Overall, with the exception of NK cells and eosinophils, which exhibited higher levels of necrosis in the graft, early apoptosis was the predominant form of cell death observed across the analyzed cell populations. Following NMP, both graft-resident CD4⁺ and CD8⁺ T cells exhibited increased rates of early apoptosis, while CD4⁺ T cells also displayed increased late apoptosis in both the graft and the perfusate. Early apoptosis was likewise increased among renal NK cells following NMP, whereas NK cells recovered from the NMP perfusate displayed lower levels of necrosis compared with HMP. Similarly, eosinophils from the NMP graft and dendritic cells from the NMP perfusate showed reduced necrosis compared with HMP. In addition, NMP induced increased early apoptosis among graft-derived eosinophils and perfusate-derived macrophages (Figure 4C). Thus, both perfusion conditions result in distinct patterns of necrosis and early apoptosis, depending on the cell type.

**Figure 4.**
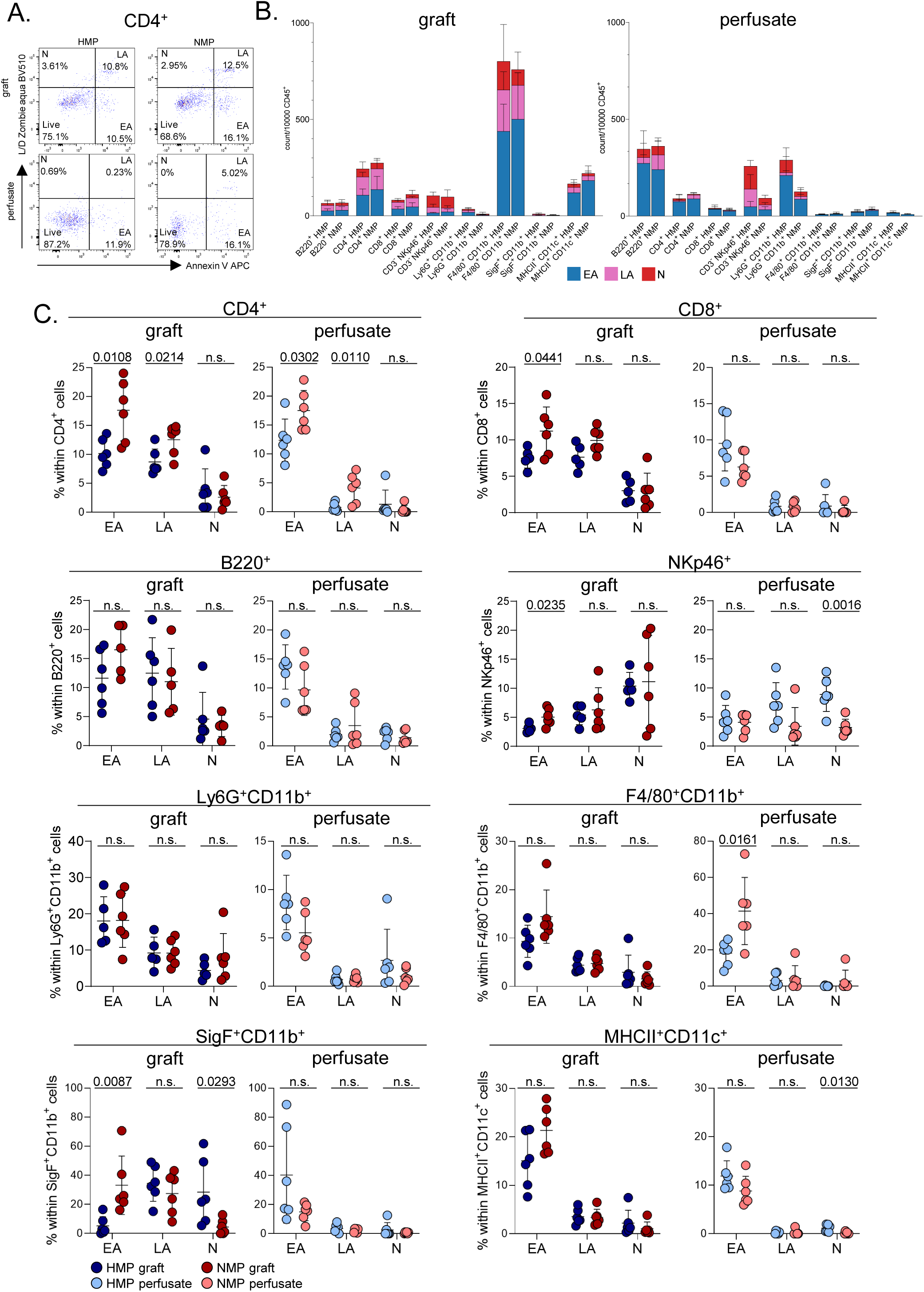
Necrosis and apoptosis in HMP- and NMP-perfused kidney grafts. (A) Representative flow cytometry dot plots showing viable (LIVE; Viability⁻/Annexin V⁻), early apoptotic (EA; Viability⁻/Annexin V⁺), late apoptotic (LA; Viability⁺/Annexin V⁺), and necrotic (N; Viability⁺/Annexin V⁻) CD4⁺ T cells isolated from the perfusate or perfused kidney grafts. FACS dot plots illustrate the respective percentage within CD4^+^ T cells. (B) Counts of early apoptotic, late apoptotic, and necrotic cells among the major leukocyte populations identified in perfusates and perfused kidney grafts. (C) Distribution of early apoptotic, late apoptotic, and necrotic cells among the major leukocyte populations identified in perfusates and perfused kidney grafts. Individual data points are shown with mean ± SD; n=5-6/experimental group. Data distribution was assessed using the Kolmogorov–Smirnov test. Statistical significance was determined using Welch’s *t*-test or Mann–Whitney *U* test, as appropriate.

### NMP enhances endothelial cell and PTECs activation

To assess the effects of HMP and NMP on renal structural cells relative to naïve kidneys and those preserved by SCS, endothelial cell activation was examined by quantifying the surface expression of the inflammatory adhesion molecules CD44^18^ and intercellular adhesion molecule-2 (ICAM-2)^19,20^. SCS markedly increased CD44 expression on CD31⁺ endothelial cells compared with naïve tissue, whereas HMP attenuated this response. In contrast, CD44 levels were significantly higher in NMP-perfused grafts than in HMP-treated kidneys. Interestingly, HMP-preserved kidneys also exhibited lower CD44 expression than naïve controls (Figure 5A). A similar pattern was observed for ICAM-2, with the highest expression detected in SCS-preserved kidneys; however, the difference between the HMP and NMP groups did not reach statistical significance (Figure 5A). Immunofluorescence staining further revealed upregulation of CD31 and ICAM-2 in both SCS-preserved and perfused kidneys (Figure 5B). Despite both perfusion strategies being associated with higher rates of late apoptosis in CD31⁺ endothelial cells than SCS, they substantially reduced the proportion of necrotic endothelial cells relative to SCS-preserved grafts (Figure 5C, D).

**Figure 5.**
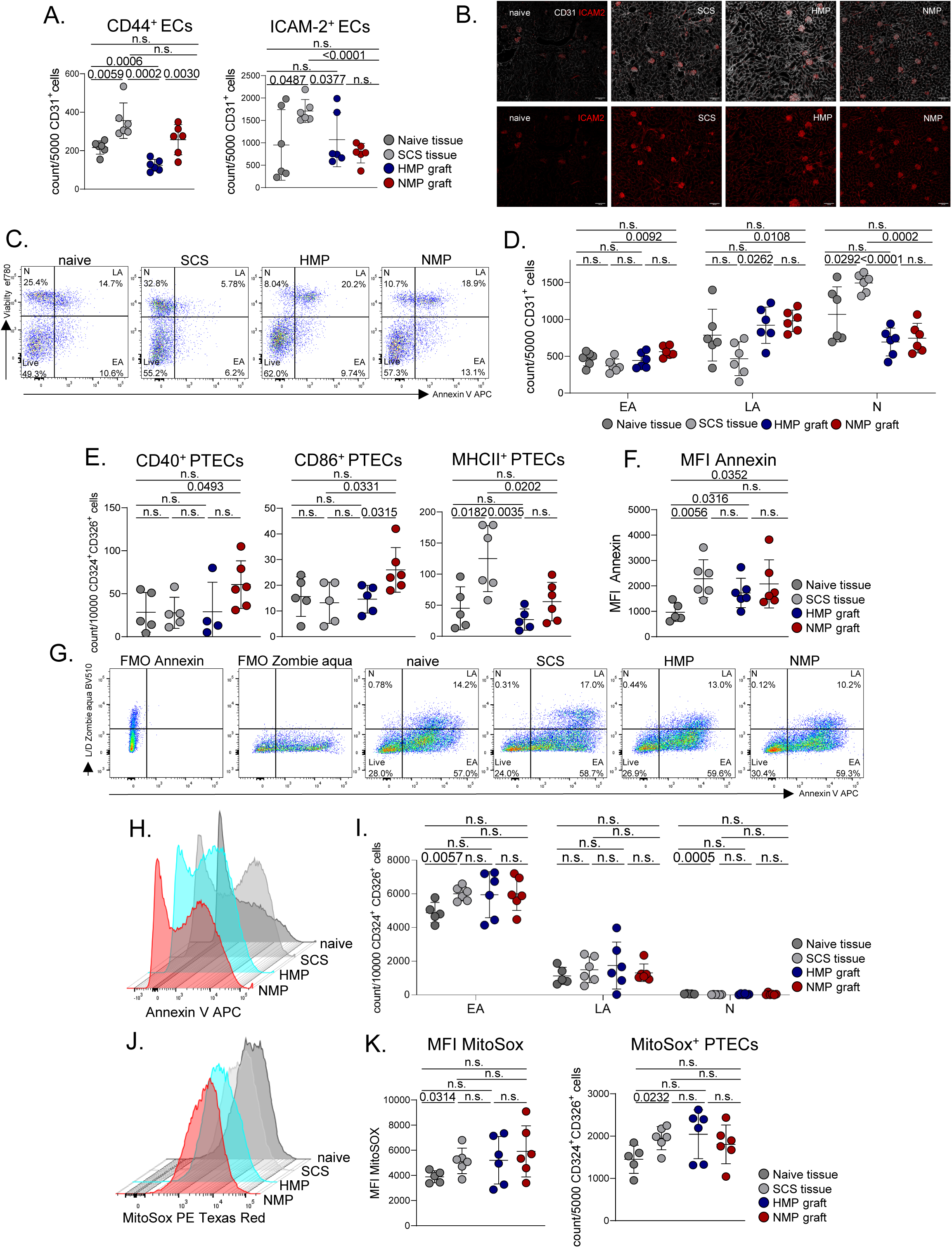
Intragraft activation of endothelial cells (ECs) and proximal tubular epithelial cells (PTECs). (A) Counts of CD31⁺CD44⁺ and CD31⁺ICAM-2⁺ ECs isolated from naïve grafts and grafts subjected to SCS, HMP, or NMP. (B) Representative immunofluorescence microscopy images of naïve kidneys and samples after SCS, HMP and NMP stained for CD31 and ICAM-2. (C) Representative flow cytometry dot plots showing viable (LIVE; Viability⁻/Annexin V⁻), early apoptotic (EA; Viability⁻/Annexin V⁺), late apoptotic (LA; Viability⁺/Annexin V⁺), and necrotic (N; Viability⁺/Annexin V⁻) CD31⁺ ECs from each experimental group. Data shown is the respective percentage within CD31^+^ ECs. (D) Absolute numbers of EA, LA, and necrotic CD31⁺ ECs from each experimental group. (E) Absolute numbers of CD40⁺, CD86⁺ and MHCII^+^ CD324⁺CD326⁺ PTECs isolated from naïve grafts and from grafts following SCS, HMP, or NMP. (F) The mean fluorescence intensity (MFI) of Annexin V was measured on PTECs from each experimental group. (G) Fluorescence minus one (FMO) controls for Annexin V and L/D Zombie aqua (Viability) and representative flow cytometry dot plots showing viable, EA, LA, and necrotic PTECs isolated from each experimental group. (H) Histogram plot of cell counts (y-axis) versus Annexin V APC fluorescence intensity (x-axis) for each experimental group. The data shown for each group are concatenated files of all PTECs from that group. (I) Absolute numbers of EA, LA, and necrotic PTECs from each experimental group. (J) Histogram plot of cell counts (y-axis) versus MitoSOX PE Texas Red fluorescence intensity (x-axis) for naïve grafts and grafts subjected to SCS, HMP or NMP. The data shown for each group are concatenated files of all PTECs from that group. (K) MFI of MitoSOX and absolute numbers of MitoSOX^+^ PTECs for each experimental group. Individual data points are shown with mean ± SD; n=5-6/experimental group. Data distribution was assessed using the Kolmogorov–Smirnov test. Statistical significance was determined using Welch’s *t*-test or Mann–Whitney *U* test, as appropriate (B,E,F,K) or using ordinary one-way ANOVA with Tukey’s multiple comparisons test (D,I).

To assess PTEC activation, the expression levels of the activation markers CD40, CD86, and MHCII were analyzed^21^. PTECs isolated from NMP-perfused kidneys demonstrated increased expression of the co-stimulatory molecules CD40 and CD86, whereas MHCII expression was significantly decreased compared with SCS-preserved tissue under both perfusion conditions (Figure 5E). Interestingly, Annexin V mean fluorescence intensity (MFI) was increased in PTECs from SCS-preserved kidneys compared with naïve tissue or HMP-perfused grafts indicating enhanced apoptotic activity (Figure 5F–H). This was further supported by a significantly higher number of early apoptotic PTECs in SCS grafts (Figure 5G). In contrast to endothelial cells, necrosis was less prominent than apoptosis in PTECs across all groups (Figure 5H, I). Finally, mitochondrial stress was assessed using MitoSOX staining. Although PTECs from both HMP- and NMP-perfused grafts showed a trend toward increased mitochondrial superoxide production, reflected by higher mean fluorescence intensity and a greater proportion of MitoSOX-positive cells, these differences did not reach statistical significance compared with naïve tissue (Figure 5J, K). Collectively, both perfusion strategies are associated with reduced inflammatory activation and a more favorable cell-death profile compared with SCS. Rather than uniformly reducing cellular injury, HMP and NMP distinctly shift the mode and distribution of injury of renal structural cells.

### Intragraft transcriptional stress-response profiles during HMP and NMP

To assess the intragraft transcriptional stress-response, we analyzed the expression of genes associated with major metabolic intracellular pathways including anaerobic glycolysis (*Slc2a1*, *glucose transporter 1* [GLUT1]; *Aldoc*, *aldolase C*), inflammation (*IL1b*, *interleukin-1β* [IL-1β]; *Ptgs2, prostaglandin-endoperoxide synthase 2* [COX-2]), lipid peroxidation (*Acsl4*, *acyl-CoA synthetase long-chain family member 4*; *Alox5*, *arachidonate 5-lipoxygenase*), iron metabolism (*Tfrc*, *transferrin receptor*), nitro-oxidative stress (*Nos2*, *inducible nitric oxide synthase* [iNOS]), antioxidant defense (*Hmox1*, *heme oxygenase 1*; *G6pdx*, *glucose-6-phosphate dehydrogenase*; *Gpx4*, *glutathione peroxidase 4*; *Keap1*, *Kelch-like ECH-associated protein 1*), hypoxia (*Egln3*, *Egl-9 family hypoxia-inducible factor 3*; *Hif1a*, *hypoxia-inducible factor 1α*), and DNA damage (*Cdkn1a*, *cyclin-dependent kinase inhibitor 1A*) (Figure 6A). *Slc2a1* expression was significantly increased following NMP, consistent with transcriptional activation of glucose-uptake and glycolysis-related pathways (Figure 6B). In addition, NMP markedly upregulated the inflammatory genes *Il1b* and *Ptgs2* compared with naïve, SCS, and HMP-treated kidneys (Figure 6C). Whereas *Acsl4* expression did not differ among groups, *Alox5* showed a trend toward higher gene expression following HMP and NMP. *Tfrc* expression was elevated across all preserved kidneys, regardless of SCS or MP, suggesting altered regulation of cellular iron uptake during organ preservation (Figure 6D, E). SCS increased *Nos2* expression, whereas this response was attenuated under HMP and NMP (Figure 6E). Accordingly, machine perfusion restored the expression of antioxidant defense genes, including *Hmox1*, *G6pdx*, and *Keap1*, and normalized the hypoxia-associated marker *Egln3 (Figure 6F).* MP also restored the expression of *Hif1a* which can be activated by osmotic stress or inflammation (Figure 6G). Despite these protective effects, NMP uniquely enhanced the DNA damage response, as indicated by significantly increased *Cdkn1a* expression (Figure 6H). Taken together, kidney grafts after oxygenated MP show transcriptional profiles associated with lower oxidative stress and enhanced antioxidant defense mechanisms, whereas NMP samples show transcriptional profiles associated with increased intragraft inflammation compared with HMP.

**Figure 6.**
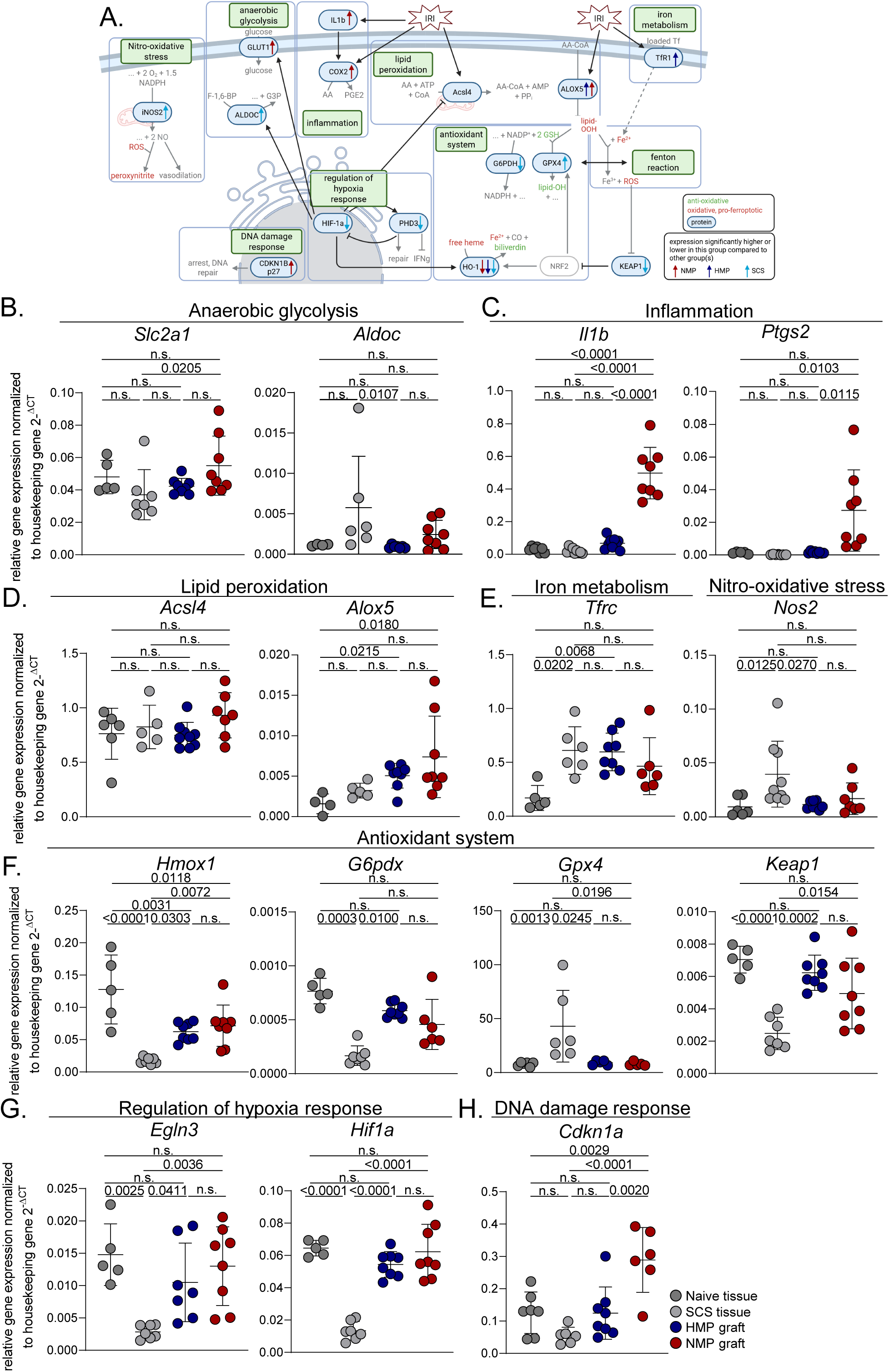
Intra-graft transcriptional stress-response profiles during HMP and NMP. (A) Schematic overview of the major intracellular pathways investigated in this study. (B–H) Gene expression analysis of genes involved in (B) anaerobic glycolysis, (C) inflammation, (D) lipid peroxidation, (E) iron metabolism and oxidative stress, (F) antioxidant defense, (G) hypoxia response, and (H) DNA damage. Individual data points are shown with mean ± SD; n=4-8/experimental group. Data distribution was assessed using the Kolmogorov–Smirnov test. Statistical significance was determined using ordinary one-way ANOVA with Tukey’s multiple comparisons test or Kruskal-Wallis test, as appropriate.

### Graft treatment with senolytics during machine perfusion

Having characterized the effects of HMP and NMP on graft-resident leukocytes and structural cell populations, we wanted to assess whether our model and analyses could be applied to test and evaluate possible graft-directed treatment strategies. Given the increasing use of expanded criteria donor (ECD) kidneys, improving the preservation of grafts from older donors with associated comorbidities is of particular interest. We previously showed that pretreatment of aged donor kidneys with the senolytic agent ABT-263 (Navitoclax), a BCL-2 family inhibitor, improves post-transplant graft quality^22^. We therefore investigated whether administration of ABT-263 during MP affects the cellular composition and integrity of the graft.

Kidneys from aged murine donors (>85 weeks) were subjected to 1 hour of oxygenated NMP supplemented with ABT-263 (Figure 7A). With the exception of CD8⁺ T cells, leukocyte frequencies in the graft were comparable to those observed in the previous experiments (Figure 7B, Figure 4B). Although the differences did not reach statistical significance, early apoptosis showed a consistent trend across the major leukocyte populations, as exemplarily shown for CD4⁺ and CD8⁺ T cells, B cells, macrophages, and dendritic cells (Figures 7C–F). ICAM-2 expression on ECs was significantly reduced following ABT-263 treatment and was accompanied by a lower frequency of CD31⁺ late-apoptotic cells within the graft (Figures 7G– I). ABT-263 did not detectably affect PTEC activation or apoptosis (Figures 7J-L). However, ABT-263-treated kidneys exhibited increased frequencies of MitoSOX⁺ PTECs, consistent with enhanced mitochondrial ROS production (Figures 7M–O). Thus, ABT-263 treatment elicited cell-type-specific alterations in apoptosis-related readouts, including increased mitochondrial oxidative stress in PTECs. Whether these changes influence allograft quality and long-term graft integrity warrants further investigation.

**Figure 7.**
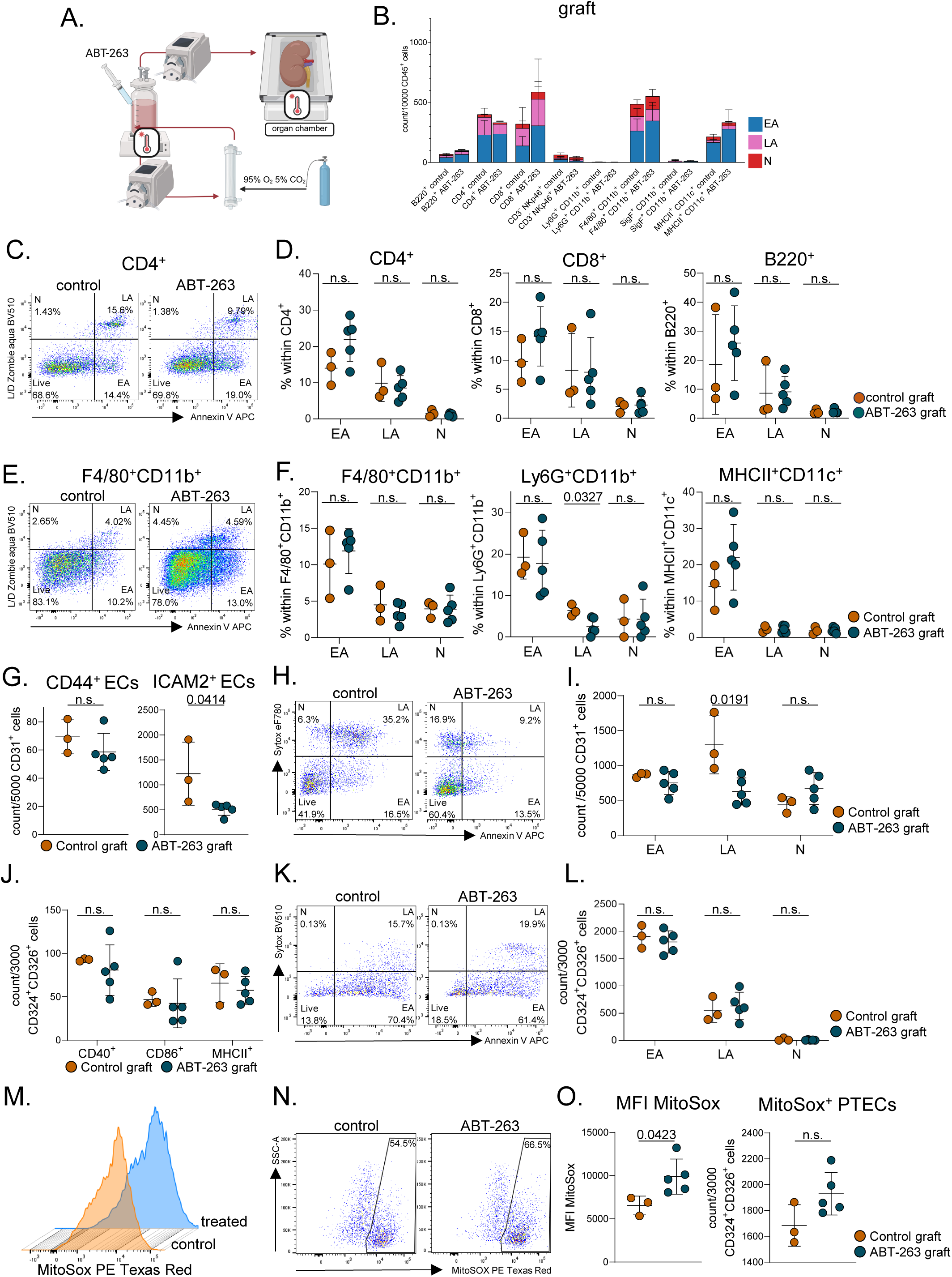
Pilot study assessing the effect of ABT-263 on kidney grafts during NMP. (A) Schematic overview of the experimental setup. (B) Counts of early and late apoptotic, and necrotic cells among the major leukocyte populations identified in perfused kidney grafts of the control and ABT-263 treatment groups. (C) Representative flow cytometry dot plots showing viable (LIVE; Viability⁻/Annexin V⁻), early apoptotic (EA; Viability⁻/Annexin V⁺), late apoptotic (LA; Viability⁺/Annexin V⁺), and necrotic (N; Viability⁺/Annexin V⁻) CD4⁺ T cells from each experimental group. Data are percentages within CD4^+^ T cells. (D) Distribution of EA, LA and necrotic cells among CD4^+^ and CD8^+^ T cells and B220^+^ B cells identified in grafts of each experimental group. (E) Representative flow cytometry dot plots showing live, EA, LA and necrotic F4/80^+^CD11b^+^ macrophages from each experimental group. Data are percentages within macrophages. (F) Distribution of EA, LA and necrotic cells among F4/80^+^CD11b^+^ macrophages, Ly6G^+^CD11b^+^ neutrophils and MHCII^+^CD11c^+^ DCs for each experimental group. (G) Absolute numbers of CD44⁺ and ICAM-2^+^ CD31^+^ ECs for each experimental group. (H) Representative flow cytometry dot plots showing live, EA, LA and necrotic ECs from each experimental group. Data are percentages within ECs. (I) Absolute numbers of EA, LA, and necrotic ECs from each experimental group. (J) Absolute numbers of CD40⁺, CD86⁺ and MHCII^+^ CD324⁺CD326⁺ PTECs for each experimental group. (K) Representative flow cytometry dot plots showing live, EA, LA and necrotic PTECs from each experimental group. Data are percentages within PTECs. (L) Absolute numbers of EA, LA, and necrotic PTECs from each experimental group. (M) Histogram plot of cell counts (y-axis) versus MitoSOX PE Texas Red fluorescence intensity (x-axis) for PTECs of each experimental group. For each group, all respective samples were concatenated. (N) Representative flow cytometry dot plots showing MitoSOX^+^ PTECs from each experimental group. Data are percentages within PTECs. (O) MFI of MitoSOX and absolute numbers of MitoSOX^+^ PTECs for each experimental group. Individual data points are shown with mean ± SD; n=3-5/experimental group. Data distribution was assessed using the Kolmogorov–Smirnov test. Statistical significance was determined using Welch’s *t*-test or Mann–Whitney *U* test, as appropriate.

## Discussion

Despite its increasing use for the preservation of ECD kidneys, the optimal MP strategy remains unclear. Non-oxygenated HMP has become routine clinical practice and is widely considered the standard approach for kidney preservation^23,24^. However, randomized clinical studies have not demonstrated improved graft function following prolonged oxygenated NMP or 1 h of end-ischemic NMP, despite encouraging findings from large-animal models^6,25–27^. These findings suggest that the biological effects of different MP strategies are not yet fully understood. In particular, the restoration of mitochondrial activity during NMP raises questions about the metabolic adaptations that occur within the graft^8,28^. Using a standardized small-animal model, we demonstrate that MP actively reshapes the intrarenal cellular landscape in a protocol-dependent manner and further show that the perfusion strategy differentially influences necrosis, apoptosis of immune-resident cells, and the activation of ECs and PTECs.

Persistent hemolysis during RBC-supported NMP may create a cytotoxic microenvironment that contributes to graft injury^29^. In line with this, we observed greater hemolysis with increasing RBC concentrations, accompanied by reduced creatinine clearance. Although NMP with 15% RBCs resulted in more pronounced tubular injury and mRNA expression of inflammatory markers, expression of established renal injury markers, including *Lcn2* and *Havcr1*, did not differ from that observed in SCS-or HMP-perfused grafts, consistent with the inconsistent findings reported previously^30,31^. Prolonged NMP with RBC-containing perfusates has been associated with progressive iron accumulation and phospholipid peroxidation in the kidney^32^. In our model, however, lipid peroxidation was only modestly increased, as indicated by a trend toward higher *Alox5* expression. This may be related to the shorter perfusion period used in our experimental set-up compared with the prolonged NMP protocols employed in previous studies^29,32^.

While the release of immune cells into the perfusate during MP has been described previously^16,31^, prior studies have not reported substantial changes in immune cell composition during perfusion^31^. In our comparison of HMP and NMP, the cellular composition of perfusates differed between the two perfusion modalities, indicating differential release of renal immune cell populations during *ex vivo* preservation. In general, leukocyte migration from the circulation across the endothelial barrier into the tissue is referred to as diapedesis, a multistep process that begins with transient selectin-mediated interactions, which slow circulating cells and facilitate chemokine detection. Subsequent chemokine signaling activates leukocyte integrins, promoting their binding to endothelial ligands such as ICAM-1, ICAM-2, and VCAM-1, followed by firm adhesion, endothelial surface migration, and trans-endothelial migration.^32^ In the present study, however, we observe cell mobilization in the opposite direction, from the tissue into the bloodstream. Therefore, the observed process may involve mechanisms analogous to those described for diapedesis but does not correspond to classical diapedesis as defined above. While neutrophil extravasation has been extensively characterized^33^, the mechanisms underlying B cell transmigration remain less well understood. EBV-infected B cells have been shown to disrupt and cross endothelial barriers, which may facilitate their infiltration into non-lymphoid tissues^34^. In our experiments, both B cells and neutrophils were released from the kidney during perfusion, consistent with previous observations of neutrophil release into the perfusate during NMP of liver grafts^35^. Although transcellular migration has been reported to be particularly prominent among effector memory (EM) CD8⁺ T cells^36^, our data indicate that both CD4⁺ and CD8⁺ EM T cells were predominantly retained within the tissue, whereas naïve T cells were more readily released into the perfusate. Further studies are needed to elucidate the expression of molecules on different leukocyte subsets that may facilitate their exit from tissues under varying conditions of temperature and perfusion. Since kidney-resident immune cells influence allograft outcomes through their roles in tissue homeostasis and immune responses, preferentially retaining kidney-resident macrophages and Tregs may be important for preserving and protecting the graft during perfusion. Conversely, retention of immune cell subsets associated with allograft injury or rejection could have opposing effects, including potential support of recipient B cell responses^37^. Thus, MP selectively affects immune-cell populations rather than causing uniform leukocyte release from the graft.

Compared with SCS, oxygenated MP was associated with lower endothelial activation, as indicated by reduced CD44 and ICAM-2 expression, and with less necrosis. Endothelial injury and necrosis during MP have previously been reported in liver perfusion and attributed to high perfusion pressures under non-oxygenated conditions^2^. We also detected PTEC activation, with early apoptosis representing a prominent form of cell death in these cells. However, neither PTEC activation nor apoptosis differed significantly between HMP and NMP. Apoptosis generally releases few damage-associated molecular patterns (DAMPs), while apoptotic debris promote an anti-inflammatory response through efferocytosis by intrarenal monocytes and macrophages^38^. In contrast, necrosis involves early loss of plasma membrane integrity and increased DAMP release, triggering innate and adaptive immune responses^39,40^. As MP significantly reduced necrosis in ECs, this may represent a beneficial effect on this cell population.

Senolytic agents such as Navitoclax induce apoptosis in senescent cells by inhibiting the anti-apoptotic proteins including BCL-2^41^. Since they are being investigated for targeting senescent cells in aged donor organs and for expanding the donor pool^22^, we conducted a pilot study to evaluate the effects of ABT-263 during NMP. After 1 hour of perfusion, ABT-263 affected renal cell populations to varying degrees, with a clear trend toward increased early apoptosis, reduced EC activation, and greater mitochondrial stress in PTECs illustrating the feasibility of our perfusion model and cellular readout system. One limitation of our study was the relatively short perfusion duration of one hour. This was necessitated by the addition of RBCs to the NMP perfusate, which resulted in a progressive increase in vascular resistance over time (data not shown). We therefore limited the NMP duration to one hour to ensure comparability with the HMP protocol. Furthermore, because the NMP perfusate contained RBCs, different cell-isolation strategies were required for the HMP and NMP perfusates. These differences may have resulted in variable cell-recovery efficiencies, particularly for granulocytes, and thus represent a potential confounding factor in the analysis of immune cell populations.

Overall, our small-animal MP model revealed distinct intrarenal responses to different perfusion modalities and allows the characterization of associated cellular changes. It therefore provides a useful platform for studying MP protocols and evaluating potential graft-directed therapies.

## Material and Methods

### Animals and kidney preparation

All experiments were performed using male C57BL/6J mice (Charles River, Sulzfeld, Germany) aged 8–14 or 70–85 weeks. The animals were housed under standardized conditions in a facility adhering to the ‘Principles of Laboratory Animal Care’ as outlined by the National Academy of Sciences and published by the National Institutes of Health (NIH Publication No. 86–23, revised 1985). All experimental procedures were conducted in accordance with relevant regulations and were approved by the Landesamt für Gesundheit und Soziales, Berlin, Germany (G0048-25). Mice were anesthetized with isoflurane and subjected to a modified surgical procedure as previously described^15^ to isolate an intact renal bundle comprising the left kidney, urinary tract, cannulated bladder, aorta, and inferior vena cava. In brief, all vascular branches except the renal vessels were ligated and transected to prevent leakage. Following median laparotomy, the intestine was mobilized to expose the bladder and left kidney. The right ureter was cauterized, and the urethra ligated and divided. The bladder was cannulated with low-density polyethylene tubing (LDPE; outer diameter 1.8 mm). Connective tissue surrounding the aorta and inferior vena cava was removed, and major branches, including the superior mesenteric and suprarenal arteries, were occluded and transected. The aorta and inferior vena cava were ligated proximal to the renal vessels, incised, flushed, and cannulated. The renal bundle was then recovered and transferred to the MP system.

To recover naïve kidney tissue samples, mice were anesthetized as described above. The abdominal fur was sanitized with 70% alcohol, followed by a midline thoracotomy and laparotomy. The vena cava was incised and the mice were transcardially perfused with Histidine-Tryptophan-Ketoglutarate (HTK) solution (Dr. Franz Köhler Chemie, Bensheim, Germany) until the internal organs appeared bloodless. The kidneys were then recovered and used for further analysis. SCS samples were recovered using the same procedure, placed in HTK solution and stored at 4°C for 1 hour prior to further analysis.

### Machine perfusion set-up

Both NMP and HMP were performed using a modified setup as published previously^15^. The full list of components is provided in Supplementary Table S1. Perfusion was conducted for one hour under pressure control at 80 mmHg. Before perfusion, the perfusate was oxygenated to a PaO₂ of 600 hPa using Carbogen (95% O₂, 5% CO₂). NMP was carried out at 37°C with the reservoir and organ chamber heated by a water bath, while HMP was maintained at 4°C. Perfusate temperature was continuously monitored. The perfusate composition is listed in Supplementary Table S2. Clinical-grade leukocyte-reduced RBCs were obtained from the local blood bank and were washed and centrifuged three times at 2000 × g, 4°C for 8 minutes to remove buffer. Human RBCs were added to the perfusate to achieve hematocrit levels of 15% or 30%. To assess the suitability of our model for evaluating treatment strategies, aged kidneys were perfused with the senolytic agent ABT-263 (Hycultec, Beutelsbach, Germany). A 1 M stock solution of ABT-263 was diluted in dimethyl sulfoxide (DMSO) and prepared as 1 mM working-stock aliquots, which were stored at −20 °C. Prior to initiating MP, 600 µL of the working stock was added to 300 mL of perfusate, resulting in a final ABT-263 concentration of 2 µM.

### Calculation of vascular resistance and flow

To evaluate the quality of pressure-controlled perfusion, perfusion pressure (mmHg) and perfusate flow rate (ml/min) were continuously recorded using BDAS software (HSE Harvard Apparatus, March-Hugstetten, Germany). Vascular resistance (VasRes) of the intrarenal vessels was calculated as: 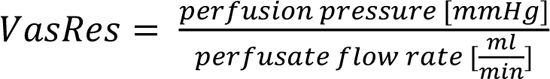

Perfusion was terminated in cases of a sudden increase in VasRes > 50 mmHg/ml/min, indicating embolism, or a sudden drop due to arterial rupture or injury.

### Hemolysis quantification

Hemolysis during NMP was assessed by measuring free hemoglobin (Hb) in venous perfusate samples via spectrophotometry at 540 nm, using a Tecan Infinite F Nano+ spectrophotometer (Tecan Life Sciences, Männedorf, Switzerland), following an established protocol^42^. A calibration curve was generated by serial dilution of a Hb standard solution (8.9 g/dl), ranging down to 8.9 × 10⁻⁹ g/dl. Absorbance readings (in duplicate) were plotted against Hb concentrations, and a linear regression yielded the equation with a goodness of fit R^2^=0.9995:

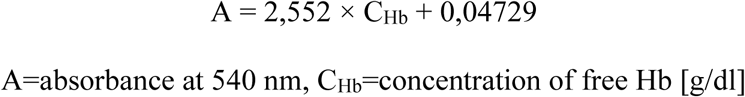

Venous perfusate samples were collected every 15 minutes, centrifuged at 1000 rpm for 10 minutes at 4°C, and 150 µl of supernatant was analyzed in a 96-well plate. Free Hb concentration was determined by applying the inverse of the regression equation to absorbance measurements.

### Assessment of kidney function: creatinine clearance and fractional sodium excretion

Kidney graft function during NMP was evaluated by calculating creatinine clearance and fractional sodium excretion (FENa⁺). Due to suppressed metabolism and lack of ultrafiltrate during HMP, kidney function assessment was limited to the NMP group. Creatinine and sodium concentrations were measured in venous perfusate and urine samples at 30 and 60 minutes using standard clinical assays on a modular analyzer using ion-sensitive electrodes and photometry (Cobas 8000 ISE and c701 modules, Roche Diagnostics, Berlin, Germany). FENa^+^ for both timepoints was calculated according to the following formula:

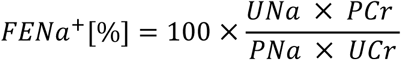

UNa = urine sodium concentration [mmol/l], PCr = perfusate creatinine concentration [mmol/l], PNa = perfusate sodium concentration [mmol/l], UCr = urine creatinine concentration [mmol/l]

Creatinine clearance was calculated using the following formula:

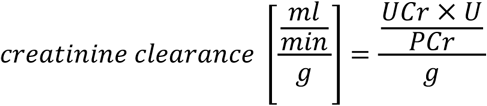

U= urine volume [ml/min], g = kidney weight [g]

### Cell isolation from perfused grafts

Renal leukocytes were isolated from kidney tissue and enriched by magnetic-activated cell sorting (MACS) using CD45 MicroBeads, as previously described^22^. Briefly, kidney tissue was mechanically dissociated and enzymatically digested for 45 min at 37 °C in a shaking water bath. The digestion mixture consisted of 10 ml RPMI 1640 medium supplemented with 10% fetal calf serum (FCS), 1% penicillin/streptomycin, collagenase IV, and DNase I. Enzymatic digestion was terminated by adding 10 ml ice-cold phosphate-buffered saline containing bovine serum albumin (PBS/BSA). The cell suspension was passed through a 100-µm cell strainer and centrifuged at 20 × g for 10 min at 4 °C to remove larger tissue fragments. The resulting supernatant was subsequently centrifuged at 300 × g for 10 min at 4 °C to pellet leukocytes. The cell pellet was resuspended in 270 µl PBS/BSA and incubated with 30 µl CD45 MicroBeads (Miltenyi Biotec, Auburn, CA, USA) for 15 min at 4 °C. After two washes with PBS/BSA, CD45-positive cells were enriched using MACS LS separation columns (Miltenyi Biotec) according to the manufacturer’s instructions.

ECs were isolated from kidney tissue using a procedure modified after Dumas^43^. Briefly, the pellet resulting from the 20 × g centrifugation step during the leukocyte isolation was used. This was resuspended in 10 ml of PBS/BSA, filtered through a 40 µm strainer and then centrifuged at 300 × g for 10 min at 4 °C. The resulting pellet contained parenchymal endothelial cells (pECs) which were used for subsequent analyses. PTECs were isolated as previously described^44^. Briefly, kidneys were cut into five slices approximately 0.5 mm thick. The tissue slices were suspended in 1 ml digestion buffer consisting of RPMI 1640 supplemented with 10% FCS, 1.25 µl DNase I, and 3.33 µl collagenase II. Samples were digested for 7 min at 37 °C in a shaking water bath. The suspension was gently pipetted up and down, after which 1 ml of the supernatant was discarded to remove tubules damaged during tissue slicing. The tissue slices were then subjected to a second digestion in 1 ml RPMI 1640 supplemented with 10% FCS and 1.25 µl DNase I for an additional 7 min at 37 °C in a shaking water bath. Following gentle pipetting, 1 ml of the resulting cell suspension was transferred to 10 ml storage solution containing RPMI 1640, 12.5 µl DNase I, and 5 mg BSA. To increase cell yield, both digestion steps were repeated, and an additional 1 ml of the resulting cell suspension was pooled with the previously collected fraction in the same storage solution. The combined cell suspension was passed through a 40-µm cell strainer, which was subsequently washed with 15 ml PBS. Cells retained on the strainer were recovered by gently scraping the filter surface with the plunger of a syringe, followed by an additional wash with 20 ml PBS. The cell suspension was centrifuged at 300 × g for 10 min at 4 °C, and the resulting cell pellet was resuspended in 3 ml PBS/BSA for downstream analyses.

### Cell isolation from MP perfusates

The entire perfusate volume of about 180 ml, depending on the flow rate, was processed for cell isolation. Peripheral mononuclear cells (PBMCs) from HMP perfusates were isolated by centrifugation at 600 × g for 12 min at 4 °C. The resulting cell pellet was collected and directly used for subsequent downstream analyses. Due to the high erythrocyte content of NMP perfusates, T cells were enriched by density gradient centrifugation using human Pancoll (PAN-Biotech, Aidenbach, Germany). Briefly, the perfusate was carefully layered onto Pancoll at a 4:1 (perfusate:Pancoll) ratio and centrifuged at 872 × g for 20 min at room temperature without brake. Mononuclear cells localized to the interphase, which was carefully collected and washed twice with PBS/BSA by centrifugation at 504 × g for 10 min at 4 °C. As granulocytes sediment together with the erythrocyte fraction during density gradient centrifugation, an alternative isolation procedure was required for the analysis of these cell populations. Residual erythrocytes were removed by incubating the cell suspension with ammonium-chloride-potassium (ACK) lysis buffer at a 1:10 dilution for 5 min at room temperature. Lysis was subsequently terminated by diluting the suspension 1:1 with PBS/BSA, followed by centrifugation at 600 × g for 12 min at 4 °C. After removal of the supernatant, the ACK lysis and washing steps were repeated once. The resulting cell pellet was then used for downstream analyses.

### Flow Cytometry and data acquisition

Viable cells were quantified using a CASY cell counter (Omni Life Science, Bremen, Germany), and typically 1 × 10⁶ cells were used per staining reaction. Freshly isolated cells were incubated with 5 µl Fc receptor blocking reagent (Miltenyi Biotec) in 45 µl PBS per sample for 10 min at 4 °C. For immune phenotyping, cells were incubated with the respective antibodies for 20 min at room temperature in the dark. For intracellular staining, cells were fixed and permeabilized using the eBioscience™ Fixation/Permeabilization Kit (Thermo Fisher Scientific, Darmstadt, Germany) according to the manufacturer’s instructions before incubation with the respective antibodies for 25 min at 37 °C in the dark. An overview of the antibodies used is provided in Supplementary Tables S3 and S4. Following incubation, samples were washed at 500 × g for 6 min at 4 °C and subsequently analyzed using a BD LSRFortessa™ X-20 flow cytometer (BD Biosciences, Heidelberg, Germany). Immune cell subsets and CD31⁺ ECs, stained with the antibodies listed in Supplementary Tables S3-5, were identified according to the gating strategies shown in Supplementary Figures S1–3A–C. To assess apoptosis and necrosis, cells were first stained as described above and subsequently incubated with 0.1 µL Viability Zombie aqua (Biolegend, San Diego, USA; 1:1000 dilution) or Viability eF780 (Thermo Fisher Scientific; 1:1000 dilution) for 10 min at room temperature. Cells were then centrifuged at 500 × g for 6 min at 4 °C, the supernatant was discarded, and the cell pellet was incubated with Annexin V (BioLegend; 1:1600 dilution) for 15 min at room temperature in the dark. Following a final washing step, cells were immediately analyzed by flow cytometry. Live cells were defined as Annexin V⁻/Viability⁻, early apoptotic cells as Annexin V⁺/Viability⁻, late apoptotic cells as Annexin V⁺/Viability⁺, and necrotic cells as Annexin V⁻/Viability⁺ ^45^. Fluorescence minus one (FMO) controls were used to define gating thresholds, as shown in Supplementary Figure S3D.

PTECs isolated from kidney tissue were identified as live, single CD45⁻CD324⁺CD326⁺ cells as shown in Supplementary Figure S4A. To assess mitochondrial oxidative stress in PTECs, cells were stained with the antibody panel listed in Supplementary Table S5 together with MitoSOX™ Red mitochondrial superoxide indicator (Thermo Fisher Scientific). Following surface staining, cells were centrifuged at 500 × g for 6 min at 4 °C and resuspended in 100 µl Hanks’ Balanced Salt Solution (HBSS) containing 0.05 µl MitoSOX™ Red. Cells were incubated for 10 min at 37 °C in the dark, washed with PBS, centrifuged again at 500 × g for 6 min at 4 °C, and immediately analyzed by flow cytometry. The MitoSOX-positive gate as well as CD44 and ICAM-2 surface expression was defined using FMO controls and unstained controls, as shown in Supplementary Figure S4B-F.

### Real-Time RT-PCR

Gene expression levels were quantified using real-time reverse transcription quantitative PCR (RT-PCR). Total RNA was isolated from snap-frozen kidney tissue using the RNeasy Mini Kit (Qiagen, Hilden, Germany) according to the manufacturer’s instructions. RNA concentration and purity were determined using a NanoPhotometer N50 (Implen, Munich, Germany). For genes assessed using commercially available TaqMan assays, cDNA was synthesized from total RNA using M-MLV reverse transcriptase and random primers. Quantitative PCR reactions were performed in a final volume of 12.75 µl, containing 1 µl cDNA, 6.25 µl TaqMan Master Mix (Thermo Fisher Scientific), 1 µl of the respective TaqMan Gene Expression Assay (Supplementary Table S6), and 5 µl nuclease-free water. Amplification was performed on a QuantStudio™ 3 Real-Time PCR System (Thermo Fisher Scientific). All samples were analyzed in duplicates. Relative expression levels of target genes were calculated using the ΔCt method and normalized to the gene expression of hypoxanthine phosphoribosyltransferase 1 (*HPRT1*). For selected target genes, custom-designed primer pairs were used (Supplementary Table S6). In these experiments, cDNA synthesis was performed using 2 µl RNA, SuperScript™ II Reverse Transcriptase (Thermo Fisher Scientific), and random hexamer primers according to the manufacturer’s protocol. Quantitative PCR was carried out using Power SYBR™ Green PCR Master Mix (Thermo Fisher Scientific) on a CFX Connect™ Real-Time PCR Detection System (Bio-Rad Laboratories, Hercules, USA). Primer sequences were designed using Primer-BLAST and synthesized by TIB Molbiol Syntheselabor GmbH, Berlin, Germany. Relative gene expression was determined using the ΔCt method and normalized to *β-actin* gene expression.

### Histology

Kidney tissues were formalin-fixed, paraffin-embedded, and sectioned at a thickness of 1 µm. Sections were processed and stained with Periodic Acid-Schiff (PAS) and counterstained with hematoxylin, as previously described^46^. Histological evaluation included assessment of necrosis, acute tubular injury, interstitial inflammation, and glomerulitis. Semiquantitative scoring criteria were adapted in accordance with the Banff classification, with tailored scales defined for each lesion type. The following scoring systems were applied: acute tubular injury: 0 = none, 1 = mild, 2 = moderate, 3 = severe; necrosis: 0 = none, 1 = 1–10% of tissue affected, 2 = 11–20%, 3 = >20%; interstitial inflammation in non-fibrotic cortex: 0 = <10%, 1 = 11–25%, 2 = 26–50%, 3 = >50%; glomerulitis: 0 = absent, 1 = present in ≥1 glomerulus. All samples were scored by an experienced pathologist in a blinded manner based on the evaluation of the entire tissue section per slide.

### Confocal microscopy

Immunofluorescence microscopy (IF) was performed as previously described^22^. Briefly, kidneys were fixed in periodate-lysine-paraformaldehyde (PLP) fixative at 4°C overnight, cryoprotected in 30% sucrose, and embedded in OCT (Roth, Karlsruhe, Germany) compound. Cryosections (16 μm) were stained overnight at room temperature with either directly conjugated or purified primary antibodies (Supplementary Table S7). The following day, sections were incubated for 1 h with the appropriate secondary antibodies, counterstained with Hoechst, and mounted using Fluoromount-G (Thermo Fisher Scientific, Darmstadt, Germany). Images were acquired using a Leica Stellaris 8 confocal microscope equipped with a white-light laser. Tile scans were obtained with a 20× objective at a resolution of 1024 × 1024 pixels and stitched using Leica LAS X software. Image processing and analysis were performed using ImageJ (v1.54r).

### Statistical Analysis

Flow cytometry data were analyzed using FlowJo v10 (BD Biosciences). If specified, datasets were downsampled using the FlowJo DownSample plugin to enable comparative analysis of subset frequencies. Dimensionality reduction was performed using opt-SNE implemented in OMIQ software (Dotmatics, Bishop’s Stortford, UK). Statistical analyses and graphical visualization were conducted using GraphPad Prism 8 (GraphPad, La Jolla, USA). Data distribution was assessed using the Kolmogorov–Smirnov test. For comparisons between two groups, Welch’s *t*-test or the Mann–Whitney *U* test was applied, as appropriate. Comparisons between multiple groups were performed using either Kruskal–Wallis test followed by Dunn’s multiple comparisons test or one-way/two-way analysis of variance (ANOVA) followed by Tukey’s multiple comparisons test, as appropriate. Unless specified otherwise, data are presented as individual data points with mean ± standard deviation (SD), with each data point representing an individual biological replicate. A *p*-value <0.05 was considered statistically significant.

## Author Contributions

JS and KK designed experiments. JS, DMK, RZ, YY, JW, RC, RL and MF performed the experiments. KH, EGS, RB and PB performed histological analyses. JS and GHP analyzed the data and designed graphs & figures. JS and KK drafted and revised the manuscript. GHP, VP and KK supervised the project. All authors approved the final version of the manuscript.

## Data availability statement

The datasets generated during this study are available from the corresponding author on reasonable request.

