## Supplementary Figures and Tables for "Hypothermic and Normothermic Machine Perfusion Differentially Shape the Renal Cellular Landscape"

### Supplementary Figure S1

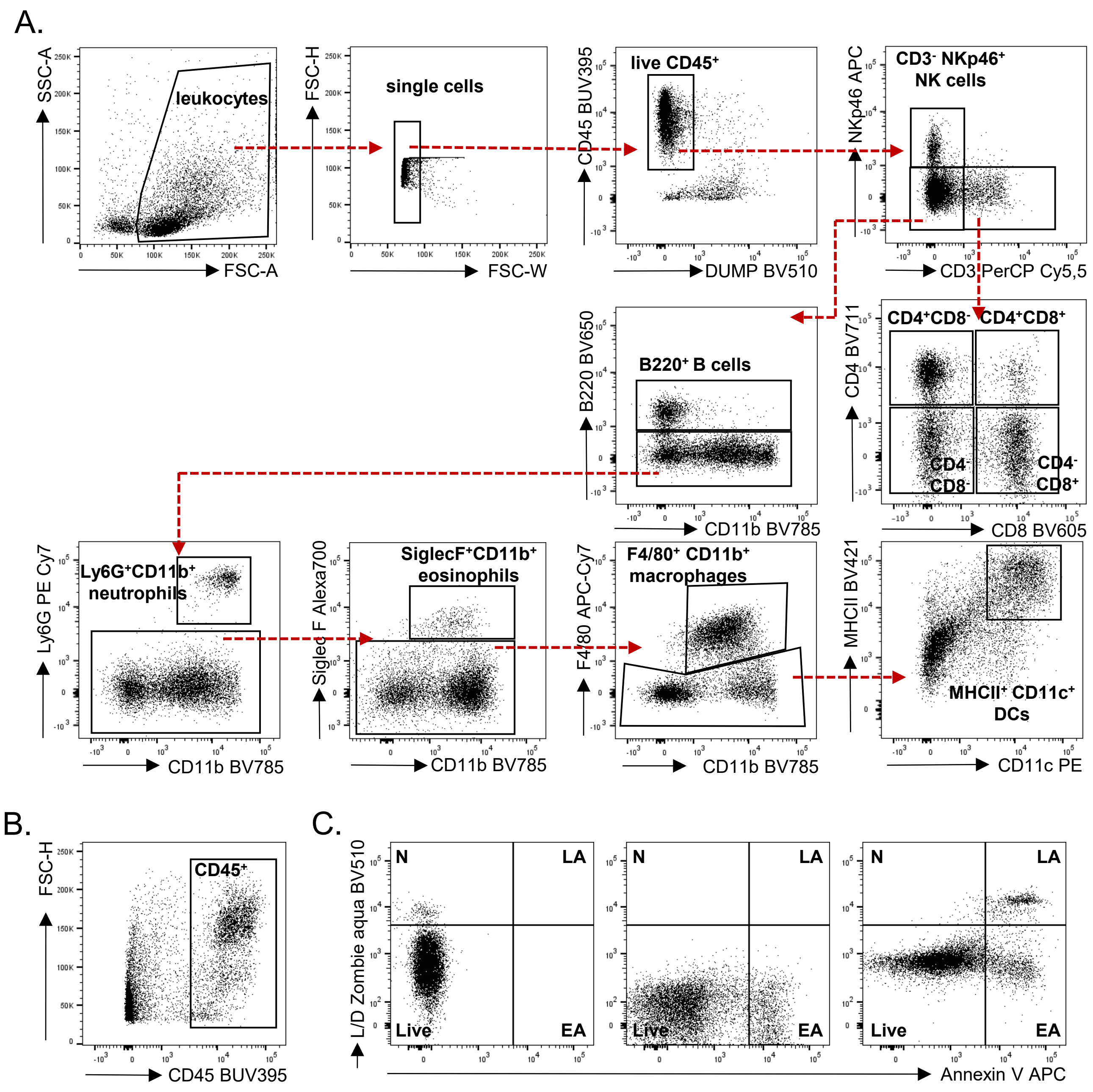


**Supplementary Figure S1. Gating strategy for the identification of immune cell subsets.** (A) Following gating on live, $CD45^{+}$ single cells, immune cell populations were further classified based on the indicated markers and gating strategy. (B) For the analysis of apoptosis and necrosis, the initial gate was set on all $CD45^{+}$ cells rather than on live $CD45^{+}$ cells. The gating strategy shown in (A) was then applied to identify the respective immune cell populations. (C) Cells were classified as live (Zombie$^{-}$ Annexin V$^{-}$), early apoptotic (EA; Zombie$^{-}$ Annexin V$^{+}$), late apoptotic (LA; Zombie$^{+}$ Annexin V$^{+}$), or necrotic (N; Zombie$^{+}$ Annexin V$^{-}$). Gates were defined based on fluorescence-minus-one (FMO) controls.

#### Supplementary Figure S2

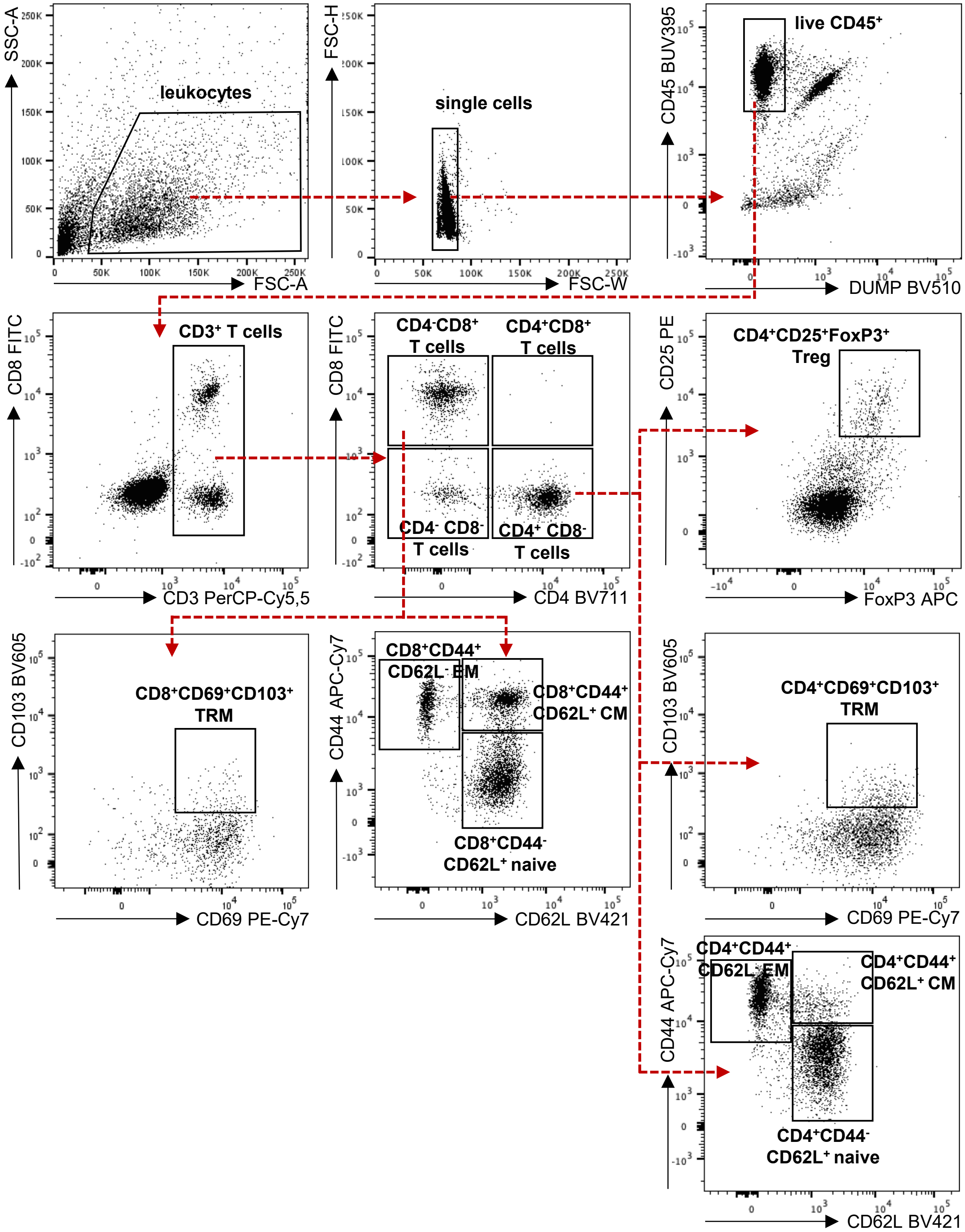


**Supplementary Figure S2.** Gating strategy for the identification of T cell subsets. Live, $CD45^{+}CD3^{+}$ single cells were identified and classified based on CD4 and CD8 expression. T cell populations were further subdivided according to the indicated markers to identify regulatory T (Treg), tissue-resident memory ($T_{RM}$), and memory T cell subsets.

#### Supplementary Figure S3

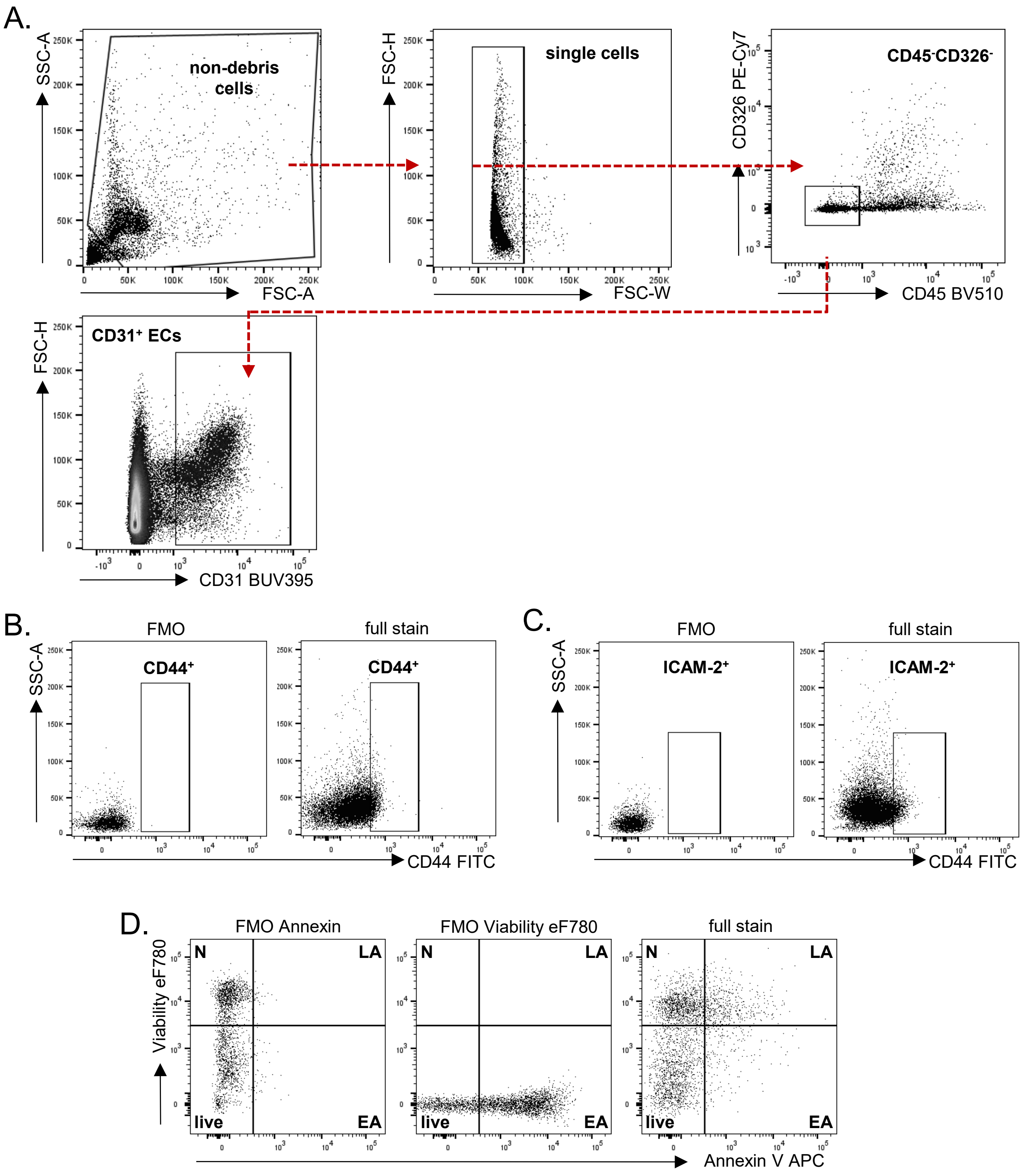


**Supplementary Figure S3.** Gating strategy for the identification of endothelial cells (ECs). (A) Endothelial cells were defined as $CD31^{+}CD45^{-}CD326^{-}$ single cells. (B) Fluorescence-minus-one (FMO) control for CD44. (C) FMO control for ICAM-2. (D) Cells were classified as live (Viability$^{-}$ Annexin V$^{-}$), early apoptotic (EA; Viability$^{-}$ Annexin V$^{+}$), late apoptotic (LA; Viability$^{+}$ Annexin V$^{+}$), or necrotic (N; Viability$^{+}$ Annexin V$^{-}$). Gates were defined based on FMO controls.

### Supplementary Figure S4

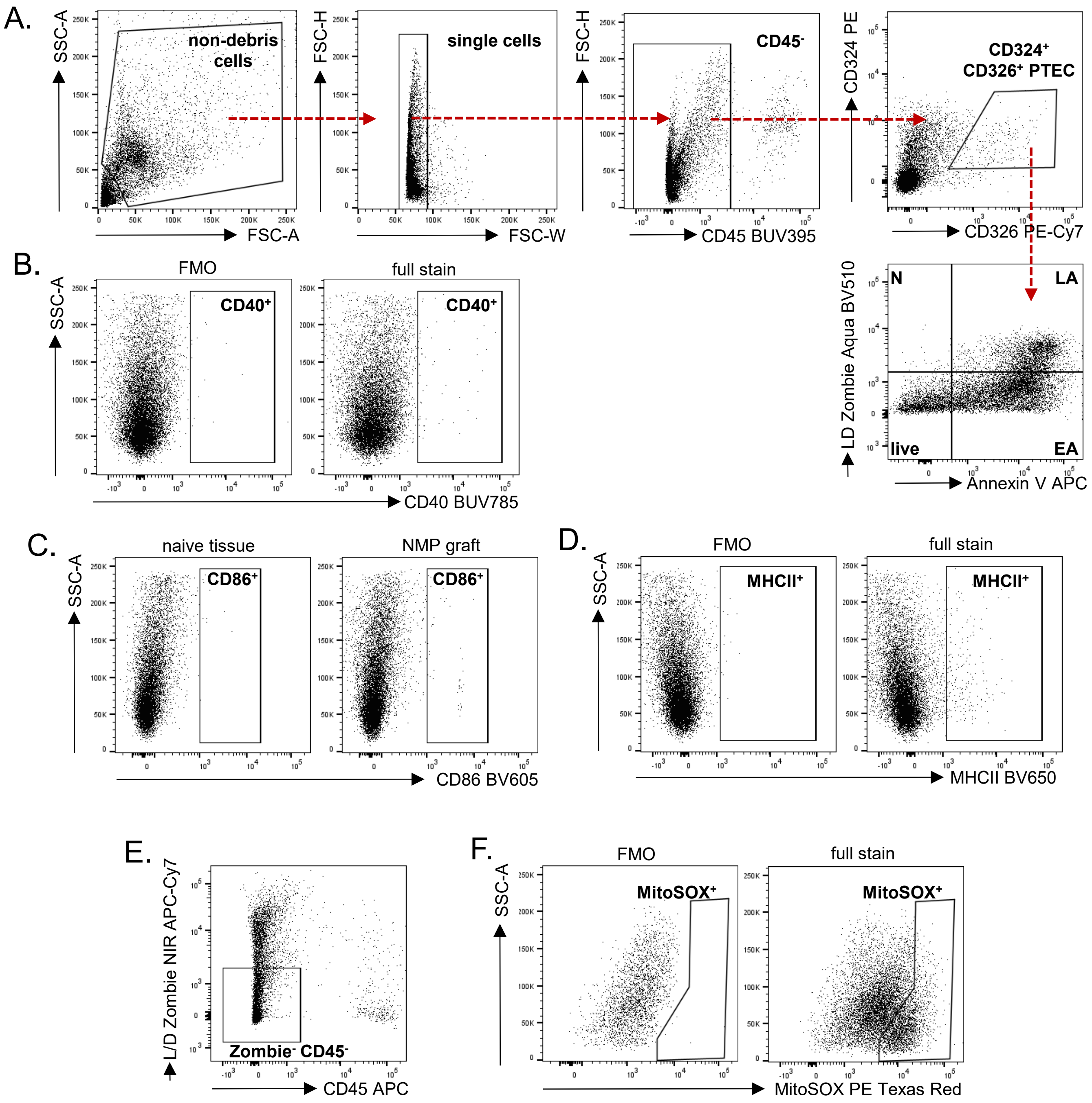


**Supplementary Figure S4**. Characterization of proximal tubular epithelial cells (PTECs). (A) Gating strategy used for the identification and characterization of PTECs). (B–D) Fluorescence-minus-one (FMO) controls for CD40, CD86, and MHC class II (MHCII), respectively. (E) For the analysis of MitoSOX positivity, the initial gate was set on viable (Zombie⁻) CD45⁻ cells. The gating strategy shown in (A) was subsequently applied to identify the PTEC population. (F) The MitoSOX-positive gate was defined based on an FMO control.

**Supplementary Table S1. Equipment used for the machine perfusion of murine kidneys.**

| Equipment | Manufacturer |
|---|---|
| Basic Data Acquisition Software (BDAS) | HSE Harvard Apparatus, March-Hugstetten, Germany |
| Bottle-Top Filter with SFCA-membrane, 0.2 µm | Carl Roth, Karlsruhe, Germany |
| Carbogen gas LAB; 95% O2, 5% CO2 | Linde Gas, Pullach, Germany |
| Glass bottle 500 ml | Brand Scientific, Wertheim, Germany |
| Glass vessels for centrifugation, 25 ml | Carl Roth, Karlsruhe, Germany |
| Grant Thermostat Optima T100 | Grant Scientific, Amsterdam, Netherlands |
| IV catheter Introcan Safety, 24G yellow, PUR, 0.7*19 mm | B. Braun Melsungen AG, Melsungen, Germany |
| Jacketed glass buffer reservoir, 1 L | HSE Harvard Apparatus, March-Hugstetten, Germany |
| Laptop HP Elite Book | Hewlett-Packard, Palo Alto, USA |
| Magnetic stirrer "Stir SB161" | Sigma Aldrich / Merck, Darmstadt, Germany |
| Masterflex Reglo digital pump, 2 channels | HSE Harvard Apparatus, March-Hugstetten, Germany |
| Millex-GP sterile filter unit 0.22 µm, 33mm, PES | Merck, Darmstadt, Germany |
| Mini peristaltic pump II | HSE Harvard Apparatus, March-Hugstetten, Germany |
| Moist chamber with bubble trap | HSE Harvard Apparatus, March-Hugstetten, Germany |
| PLUGSYS casing with TAM-D, APT300 pressure sensor and SCP controller | HSE Harvard Apparatus, March-Hugstetten, Germany |
| Pt100 temperature probe | HSE Harvard Apparatus, March-Hugstetten, Germany |
| Pump tubing, AME 14 - SC0062 (Tygon) | HSE Harvard Apparatus, March-Hugstetten, Germany |
| Silicone tubing ID 1.47mm OD 1.96 mm, length: 15 m | VWR, Radnor, USA |
| Single channel fiber optic minisensor oxygen with flow-through cell | HSE Harvard Apparatus, March-Hugstetten, Germany |
| Stand with triangular plate | HSE Harvard Apparatus, March-Hugstetten, Germany |
| Transfusion system Sangofix B | B. Braun Melsungen AG, Melsungen, Germany |
| Tubing LDPE 1.8 mm OD, 0.4 mm wall thickness, 100 m | Sigma Aldrich / Merck, Darmstadt, Germany |
| Tubing Tygon LMT-55; ID 1.22 mm | HSE Harvard Apparatus, March-Hugstetten, Germany |
| Y connectors for tubing | HSE Harvard Apparatus, March-Hugstetten, Germany |

**Supplementary Table S2. Components of the perfusate.**

| Reagent | Final concentration in perfusate | Manufacturer |
|---|---|---|
| Albumin | 4 % vol. | Carl Roth, Karlsruhe, Germany |
| Aminoplasmal 10% | 9 ml/l | B. Braun Melsungen AG, Melsungen, Germany |
| Ampicillin sodium salt | 0.5 mg/ml | Carl Roth, Karlsruhe, Germany |
| CaCl | 0.935 mMol /l | Sigma Aldrich / Merck, Darmstadt, Germany |
| Creatinine | 15 mg/l | Sigma Aldrich / Merck, Darmstadt, Germany |
| Custodiol HTK solution | for flushing | Dr. Franz Köhler Chemie, Bensheim, Germany |
| Erythrocyte concentrate (human), any blood group | 0 % vol. / 15% vol. / 30 % vol. | DRK blood donation service, Berlin, Germany |
| Furosemide, 97+% | 50 mg/l | Thermo Fisher Scientific, Schwerte, Germany |
| Glucose | 3.5-5.5 mmol/l | Merck, Darmstadt, Germany |
| Heparin 25 000 IU / 5 ml | 2000 IU / l | Panpharma, Luitré-Dompierre, France |
| KCl | 330 mg/l | Merck, Darmstadt, Germany |
| Mannitol | 2,2 mg / ml | Carl Roth, Karlsruhe, Germany |
| MgCL2 * 6 H20 | 205 mg/l | Carl Roth, Karlsruhe, Germany |
| Na2HPO4 | 120 mg/l | Carl Roth, Karlsruhe, Germany |
| NaCl | 107.8 mmol/l | Carl Roth, Karlsruhe, Germany |
| NaH2PO4 *H2O | 400 mg/l | Fluka Chemika, Seelze, Germany |
| NaHCO3 | 29 mmol/l | Carl Roth, Karlsruhe, Germany |
| Prostaglandin E2 (PGE2) | 40 µg/l | Merck, Darmstadt, Germany |
| Sterile water | | Fresenius Kabi, Bad Homburg, Germany |
| Verapamil hydrochloride | for flushing, 0.25 mg / 5 ml | VWR, Radnor, USA |
| For samples in the ABT-263 treatment group: | | |
| Navitoclax ABT-263 | 2 µm | Hycultec, Beutelsbach, Germany |

**Supplementary Table S3. Antibodies for phenotyping of immune cells.**

| Antigen | Clone | Fluorochrome | Manufacturer | Catalog # |
|---|---|---|---|---|
| B220 | RA3-6B2 | BV650 | Biolegend, San Diego, USA | 103241 |
| CD11b | M1/70 | BV785 | Biolegend, San Diego, USA | 101243 |
| CD11c | N418 | PE | Biolegend, San Diego, USA | 117308 |
| CD3 | REA641 | PerCPCy5.5 | Miltenyi, Bergisch-Gladbach, Germany | 130-109-841 |
| CD4 | RM4-5 | BV711 | Biolegend, San Diego, USA | 100550 |
| CD45 | 30-F11 | BUV395 | BD, Heidelberg, Germany | 564279 |
| CD8a | 53-6.7 | BV605 | Biolegend, San Diego, USA | 100744 |
| Ep-CAM | G8.8 | PE-Cy7 | Biolegend, San Diego, USA | 118215 |
| F4/80 | BM8 | APC-Cy7 | Biolegend, San Diego, USA | 123117 |
| Ly6C | HK1.4 | PE-Texas Red | Biolegend, San Diego, USA | 128043 |
| Ly6G | 1A8 | PE-Cy7 | Biolegend, San Diego, USA | 127617 |
| MHCII | M5/114.15.2 | BV421 | Biolegend, San Diego, USA | 107631 |
| NKp46 | 29A1.4 | APC | Biolegend, San Diego, USA | 137608 |
| Siglec-F | E50-2440 | AF700 | BD, Heidelberg, Germany | 565183 |
| Viability Zombie aqua | - | aqua (BV510) | Biolegend, San Diego, USA | 423101 |
| For apoptosis & cell death assessment: | | | | |
| Annexin V kit | - | APC | Biolegend, San Diego, USA | 640919 |
| NKp46 | 29A1.4 | FITC | eBioscience, ThermoFisher, Darmstadt, Germany | 11-3351-82 |

**Supplementary Table S4. Antibodies for phenotyping of T cell subsets.**

| Antigen | Clone | Fluorochrome | Manufacturer | Catalog # |
|---|---|---|---|---|
| extracellular | | | | |
| B220 | RA3-6B2 | BV510 | Biolegend, San Diego, USA | 103248 |
| CCR6 | 29-2L17 | BV785 | Biolegend, San Diego, USA | 129823 |
| CD103 | 2E7 | BV605 | Biolegend, San Diego, USA | 121433 |
| CD25 | BC61 | PE | Biolegend, San Diego, USA | 102008 |
| CD3 | REA641 | PerCPCy5.5 | Miltenyi, Bergisch-Gladbach, Germany | 130-109-841 |
| CD38 | 90 | AF700 | ThermoFisher, Darmstadt, Germany | 56-0381-82 |
| CD4 | RM4-5 | BV711 | Biolegend, San Diego, USA | 100550 |
| CD44 | IM7 | APC-eFluor 780 | eBioscience, ThermoFisher, Darmstadt, Germany | 47-0441-82 |
| CD45 | 30-F11 | BUV395 | BD, Heidelberg, Germany | 564279 |
| CD62L | MEL-14 | BV421 | Biolegend, San Diego, USA | 104436 |
| CD69 | H1.2F3 | PE-Cy7 | Biolegend, San Diego, USA | 104512 |
| CD8a | 53-6.7 | FITC | Biolegend, San Diego, USA | 100726 |
| Viability Zombie aqua | - | aqua (BV510) | Biolegend, San Diego, USA | 423101 |
| intracellular | | | | |
| Ki67 | SolA15 | PE-Texas Red | ThermoFisher, Darmstadt, Germany | 61-5698-82 |
| FOXP3 | MF-14 | AF647 | Biolegend, San Diego, USA | 126408 |

**Supplementary Table S5. Antibodies for phenotyping of ECs, PTECs and mitochondrial stress.**

| Antigen | Clone | Fluorochrome | Manufacturer | Catalog # |
|---|---|---|---|---|
| Endothelial cells: | | | | |
| Annexin V | - | APC | Biolegend, San Diego, USA | 640919 |
| CD102 / ICAM2 | REA745 | PE | Miltenyi, Bergisch-Gladbach, Germany | 130-112-029 |
| CD31 | MEC13.3 | BUV395 | BD, Heidelberg, Germany | 740231 |
| CD44 | IM7 | FITC | BD, Heidelberg, Germany | 561859 |
| CD45 | 30-F11 | BV510 | Biolegend, San Diego, USA | 103137 |
| Ep-CAM CD326 | G8.8 | PE-Cy7 | Biolegend, San Diego, USA | 118215 |
| Viability eF780 | - | BV785 | eBioscience, ThermoFisher, Darmstadt, Germany | 65-0865-14 |
| PTECs apoptosis & cell death assessment, activation markers: | | | | |
| Annexin V | - | APC | Biolegend, San Diego, USA | 640919 |
| CD40 | Mrz 23 | BUV785 | Biolegend, San Diego, USA | 124645 |
| CD45 | 30-F11 | BUV395 | BD, Heidelberg, Germany | 564279 |
| CD86 | GL-1 | BV605 | Biolegend, San Diego, USA | 105037 |
| E-Cadherin / CD324 | DECMA-1 | PE | Biolegend, San Diego, USA | 147303 |
| Ep-CAM / CD326 | G8.8 | PE-Cy7 | Biolegend, San Diego, USA | 118215 |
| MHCII | M5/114.15.2 | BV650 | Biolegend, San Diego, USA | 107641 |
| Viability Zombie aqua | - | aqua (BV510) | Biolegend, San Diego, USA | 423101 |
| PTECs MitoSOX: | | | | |
| CD45 | 104 | APC | Biolegend, San Diego, USA | 103111 |
| E-Cadherin / CD324 | DECMA-1 | BV421 | Biolegend, San Diego, USA | 147319 |
| Ep-CAM / CD326 | G8.8 | PE-Cy7 | Biolegend, San Diego, USA | 118215 |
| L/D Zombie NIR | - | APC-Cy7 | Biolegend, San Diego, USA | 423105 |
| MitoSox Red | - | PE | ThermoFisher, Darmstadt, Germany | M26007 |

**Supplementary Table S6. List of assays used for RT-qPCR.**

| Gene name | Assay ID | Manufacturer |
|---|---|---|
| *Acsl4* | Mm00490331_m1 | Thermo Fisher Scientific, Schwerte, Germany |
| *Alox5* | Mm01182747_m1 | Thermo Fisher Scientific, Schwerte, Germany |
| *Cdkn1a* | Mm01303209_m1 | Thermo Fisher Scientific, Schwerte, Germany |
| *Gpx4* | Mm04411498_m1 | Thermo Fisher Scientific, Schwerte, Germany |
| *Il1b* | Mm00434228_m1 | Thermo Fisher Scientific, Schwerte, Germany |
| *Nos2* | Mm00440502_m1 | Thermo Fisher Scientific, Schwerte, Germany |
| *Ptgs2* | Mm00478374_m1 | Thermo Fisher Scientific, Schwerte, Germany |
| *Tfrc* | Mm00441941_m1 | Thermo Fisher Scientific, Schwerte, Germany |
| **Gene name** | **Sequence** | |
| *Actb* | Forward: CTG TCG AGT CGC GTC CAC C<br>Reverse: GGC CTC GTC ACC CAC ATA GG | |
| *Aldoc* | Forward: GCT GAG CCA AAT TGG GGT GG<br>Reverse: CCA TCC AGC CCT TGA GTG GT | |
| *Egln3* | Forward: GCT GCG AGG CCA TCA ACT TC<br>Reverse: ATG GGC TCC ACG TCT GCT AC | |
| *G6pdx* | Forward: GAA ACT GGC TGT GCG CTA<br>Reverse: GGC TCA CTC TGC TTT CGG AT | |
| *Havcr1* | Forward: TCA GGG TCT CCT TCA CAG CAG<br>Reverse: TGA CCC ACC ACC CCC TTT AC | |
| *Hif1a* | Forward: TCG GAC AGC CTC ACC AGA CA<br>Reverse: GCT TGA AAA AGG GAG CCA TCA | |
| *Hmox1* | Forward: AAC ATC GAC AGC CCC ACC AA<br>Reverse: GCT AGC AGG CCT CTG ACG AA | |
| *Keap1* | Forward: CAT GGC TCA CAA AGT GGT GC<br>Reverse: AAG AAC TCC TCC TGC TTG GC | |
| *Lcn2* | Forward: CCA GAC TTC CGG AGC GAT CA<br>Reverse: GTG GTG GCC ACT TGC ACA TT | |
| *Slc2a1* | Forward: TCT TAA GTG CGT CAG GGC GT<br>Reverse: CTC CGT AGC GGT GGT TCC AT | |

**Supplementary Table S7. List of stains and antibodies used for immunofluorescence microscopy.**

| Antigen | Clone | Fluorochrome | Manufacturer | Catalog # |
|---|---|---|---|---|
| Hoechst 33258 | | | ThermoFisher, Darmstadt, Germany | |
| ICAM2 | mlC2/4 | AF647 | Biolegend, San Diego, USA | |
| CD31 | polyclonal | | R&D Systems, Minneapolis, USA | |
| Donkey a-goat | polyclonal | CF594 | Biotium Sigma Aldrich / Merck, Darmstadt, Germany | |
